# STING suppresses cancer pain via immune and neuronal modulation

**DOI:** 10.1101/2021.01.18.426944

**Authors:** Kaiyuan Wang, Christopher R. Donnelly, Changyu Jiang, Yihan Liao, Xueshu Tao, Sangsu Bang, Michael Lee, Matthew J. Hilton, Ru-Rong Ji

## Abstract

Agonists of the innate immune regulator stimulator of interferon genes (STING) have shown great efficacy in promoting antitumor immunity in preclinical models, leading to their exploration in cancer immunotherapy trials. Patients with advanced stage cancers frequently suffer from severe pain as a result of bone metastasis and bone destruction, for which there is no efficacious treatment. Here, using multiple mouse models of metastatic bone cancer, we report that STING agonists confer remarkable protection against cancer pain, bone destruction, and local tumor burden. Repeated systemic administration of STING agonists robustly attenuated bone cancer-induced pain symptoms and improved locomotor function. Interestingly, STING agonists provided acute pain relief through direct neuronal modulation, as *ex vivo* incubation of STING agonists reduced excitability of pain-sensing nociceptive neurons from tumor-bearing mice. In addition, STING agonists protected local bone destruction and reduced local tumor burden through modulation of osteoclast and immune cell function in the tumor microenvironment, providing long-term cancer pain relief. Finally, these *in vivo* effects were dependent on host-intrinsic STING and *Ifnar1*. Overall, STING activation provides unique advantages in controlling metastatic bone cancer pain through distinct and synergistic actions on nociceptors, immune cells, and osteoclasts.

## Introduction

Patients with advanced lung, breast, thyroid and bladder cancers frequently suffer from cancer pain following bone metastasis, which is accompanied by osteolytic lesions and severe pain ^1^. Studies estimate that approximately 75% of patients with late stage cancer experience moderate or severe pain ^2–4^, and more than half of all patients with metastatic cancer pain report insufficient pain relief by the currently available pharmacotherapies ^5, 6^. Inadequate pain control for patients with metastatic cancer is often accompanied by depression, anxiety, impaired function, and significantly reduced quality of life, leading to increased morbidity and mortality ^7–9^. Thus, in addition to the ongoing challenge of developing new therapeutics capable of treating the underlying cancer, there is also an urgent unmet clinical need to develop new therapies which provide cancer patients with palliative care to relieve pain and improve quality of life. Notably, endocannabinoid has been shown to alleviate murine bone cancer pain without producing the side effects of opioids ^10^. An ideal therapeutic approach would be one that is capable of actively treating the underlying cancer while concurrently suppressing cancer-associated pain, thereby treating the pain and the underlying disease.

STING, or stimulator of interferon (IFN) genes, is an intracellular DNA sensor which plays a critical role in innate immunity, promoting the elimination of pathogens and damaged host cells via the induction of type-I IFN (IFN-I), including IFN-α and IFN-β. Activation of the STING pathway can also potently enhance antitumor immunity, underscored by preclinical studies in which the murine STING agonist DMXAA or the cross-species STING agonist ADU-S100 have been demonstrated to suppress tumor progression and increase survival in an adaptive immune cell-dependent manner ^11–16^. In addition, several groups have demonstrated STING-activating-micro- or nanoparticles also show efficacy in promoting innate and adaptive immunity in orthotopic and genetically engineered tumor models in mice ^17–19^. These studies have led to the exploration of ADU-S100 and other small molecule STING agonists to be tested as potential immunotherapy agents in several ongoing clinical trials. Our recent study has shown that STING agonists could effectively control nociception in naïve animals and animals with pathological pain^20^. However, it remains unclear if STING agonists are effective in treating bone cancer, especially given that bone marrow is regarded to be an immunosuppressive tumor microenvironment ^21^.

Previous studies have demonstrated that IFN-I signaling suppresses osteoclast formation, and activation of the STING pathway promotes bone formation in a murine bone autoimmune disease model ^22–25^. This is noteworthy, as tumor-induced over-activation of osteoclasts is a dominant mechanism leading to osteolytic bone lesions and cancer pain ^26^. It is generally believed that activation of pain-sensing nociceptive neurons (nociceptors) by soluble mediators released from cancer cells and osteoclasts drives bone cancer pain. Based on these studies, and taken in conjunction with the promise STING agonists have shown as cancer immunotherapy agents, we posited that activation of the STING pathway in bone cancer may be a unique synergistic approach to concurrently promote antitumor immunity, suppress bone destruction, and provide pain control. In this study, we tested this hypothesis using syngeneic mouse models of bone cancer pain in which lung or breast cancer cells are introduced into the intramedullary canal of murine femurs. Using this model, we administered multiple STING agonists and measured the effects of STING pathway activation on metastatic bone cancer-induced pain, bone destruction, and local tumor burden.

## Results

### STING agonists attenuate bone cancer-induced pain and restore locomotor function

We first sought to determine whether activation of STING via systemic administration of DMXAA could provide long-term therapeutic effects in a mouse model of metastatic bone cancer. To this end, we established a syngeneic murine bone cancer pain model by inoculating Lewis lung carcinoma (LLC, 2 × 10^5^ cells in 2 µl) cells into the femora of C57BL/6 mice. Vehicle or DMXAA (20 mg/kg) was intraperitoneally (i.p.) injected twice to these mice on day 3 (3d) and day 7 (7d) after tumor implantation. Behavioral tests including von Frey testing for mechanical allodynia and acetone response duration for cold allodynia were performed on the hindpaw of the tumor-bearing leg at baseline (BL), 7d (before drug injection), 10d, and 14d after LLC inoculation. Flinches and guarding behavior for spontaneous/ongoing pain was evaluated on day 14 post tumor injection (**Fig. 1a**). DMXAA treatment significantly reduced mechanical allodynia on d7, d10 and d14 (**Fig. 1b**) and cold allodynia on d10 and d14 after LLC implantation (**Fig. 1c**). On d14, DMXAA treatment also attenuated measures of spontaneous and ongoing pain (**Fig. 1d**). No apparent sex differences were observed, as the therapeutic effect of DMXAA on bone cancer pain existed in both male and female mice (**Extended Data Fig. 1a-d**). In addition, we observed no differences in body weight between the vehicle- and DMXAA-treated groups (**Fig. 1e**), indicating the experimental protocol we used is relatively safe and without gross systemic gastrointestinal (GI) toxicity. Notably, survival was not the endpoint of this study and given that animals in late stages of this model experienced severe pain and functional impairment, all mice were sacrificed at d17 post-inoculation to maintain reasonable health conditions and minimize suffering, as indicated in our recent study ^27^.

**Figure 1.**
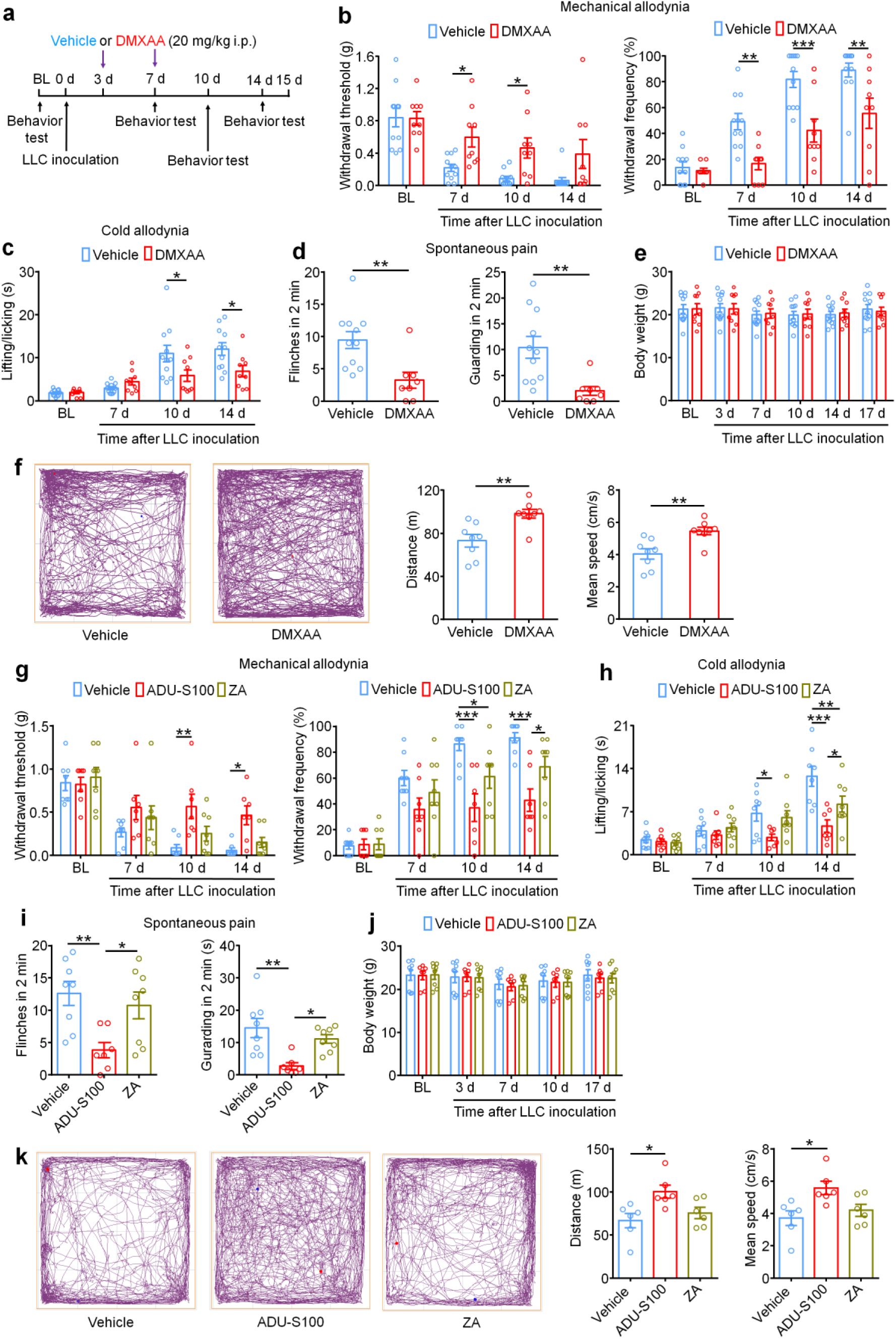
STING agonists reduce bone cancer-induced pain and functional impairment. **a.** Experimental design to test the antinociceptive effects of DMXAA in the LLC model of bone cancer. **b.** Von Frey testing to determine cancer-induced mechanical allodynia, as assessed by withdrawal threshold (left) or withdrawal frequency (0.16g stimulus; right) in mice treated with vehicle or DMXAA (20 mg/kg, i.p.) (n = 9-11 mice/group). **c.** Assessment of cancer-induced cold allodynia after LLC inoculation in mice treated with vehicle or DMXAA (n = 9-11 mice/group). **d.** Comparison of spontaneous pain as indicated by flinching behaviors (left) or guarding behaviors (right) in vehicle and DMXAA-treated mice on d14 after tumor implantation (n = 8-11 mice/group). **e.** Measurement of body weight in vehicle or DMXAA-treated mice at the indicated timepoints (n = 9 or 11). **f.** Open field testing at d14 post-inoculation to determine distance traveled (m) and mean speed (cm/s) over a 30 min duration in vehicle or DMXAA treated mice (n=8/group). Left: representative traces. Right: quantification. **g.** Von Frey testing to measure cancer-induced mechanical allodynia in mice treated with vehicle, ADU-S100 (20 mg/kg, i.p.) or ZA (100 µg/kg, i.p.), n = 7-8 mice per group. **h.** Measurement of cancer-induced cold allodynia in mice with indicated treatment on day 7, 10 and 14 after LLC inoculation (n = 7-8). **i.** Spontaneous pain as determined by flinching behaviors (left) or guarding behaviors (right) over a 2 minute interval on d14 post inoculation (n = 7-8). **j.** Body weight measurements in mice with the indicated treatments (n = 7-8). **k.** Open field testing, measuring distance traveled (m) and mean speed (cm/s) over a 30 min duration in vehicle, ADU-S100, and ZA-treated mice at d14 after tumor inoculation (n = 7-8). Left: representative traces; right: quantification. All data displayed represent the mean ± SEM. \**P* < 0.05, \*\**P* < 0.01 and \*\*\**P* < 0.001, repeated-measures two-way ANOVA with Bonferroni’s *post-hoc* test (**b, c, e, g, h, j**); two-tailed Student’s t-test (**c, e**); one-way ANOVA with Bonferroni’s *post-hoc* test (**i, k**).

Clinically, a critical comorbidity of bone metastasis in patients with advanced stage cancers is diminished or lost mobility, leading to functional impairment and reduced quality of life ^7^. To determine whether STING activation with DMXAA could improve locomotor function, we evaluated the movement activity in the open field test. Importantly, mice treated with DMXAA exhibited greater overall distance of movement and increased speed of movement on d14 after tumor inoculation (**Fig. 1f**). Thus, systemic treatment with DMXAA significantly improved locomotor function in mice with bone cancer.

Clinically, bisphosphonates are widely used for the prevention and treatment of metastatic bone cancer-induced skeletal-related events (SREs) by promoting apoptosis of bone-resorbing osteoclasts. Zoledronic acid (ZA) is one of the most potent bisphosphonates ^28^, and has also been reported to exhibit antitumor effects ^29^. Moreover, given the limited translational significance of DMXAA due to its specificity for murine STING, we also sought to test whether ADU-S100 could exert similar therapeutic effects. Following administration of vehicle, ADU-S100 (20 mg/kg), or ZA (100 µg/kg, a highly effective dose with minimal toxicity, as demonstrated in previous studies ^28, 29^ at d3 and d10, we found that ZA failed to reduce cancer-induced mechanical allodynia when analyzing paw withdrawal threshold, but could reduce withdrawal frequency to low-threshold stimulation (0.16g Von Frey filament) on d10 post tumor implantation. ADU-S100 treatment, however, could significantly attenuate mechanical allodynia in both measures on d10 and d14, with effects superior to those of ZA (**Fig. 1g**). ZA reduced cold allodynia on d14 after tumor inoculation, whereas ADU-S100 reduced cold allodynia on both d10 and d14, and this effect was significantly greater in the ADU-S100 group compared to the ZA group on d14 (**Fig. 1h**). In addition, mice treated with ADU-S100 but not ZA exhibited reduced spontaneous and ongoing pain compared to vehicle-treated mice (**Fig. 1i**). We found that these doses of ADU-S100 and ZA again had no effect on overall body weight (**Fig. 1j**), suggesting they are relatively safe and without gross GI toxicity. To again assess the potential benefits of ZA and ADU-S100 on function and mobility, we performed open field testing at d14 on mice treated with vehicle, ADU-S100 and ZA at d3 and d10. Notably, mice in the ADU-S100 treatment group but not the ZA treatment group exhibited increased overall distance of movement and increased speed of movement compared with the vehicle treatment group (**Fig. 1k**). Taken together, ADU-S100 was superior to ZA in reducing cancer-induced pain and improving locomotor function.

### STING agonists protect against cancer-induced bone destruction

Bone cancer-induced pain usually develops in tandem with the onset of tumor-induced bone destruction. It is understood that bone cancer pain is evoked by factors produced directly by cancer cells which act on afferent nociceptive nerve fibers in the tumor microenvironment (TME) ^30, 31^. In addition, cancer cells promote bone cancer pain indirectly by accelerating osteoclastogenesis, generating osteoclasts which release pro-nociceptive factors and promote bone resorption, facilitating bone destruction and painful fractures^27, 32^. Thus, cancer cells in the bone tumor microenvironment promote bone cancer through direct and indirect mechanisms (**Extended Data Fig. 1e**. *In vivo,* the LLC cell line is known to induce osteolytic bone destruction due to tumor-induced activation of osteoclast formation and activity^27^, recapitulating the pathogenesis of metastatic bone cancers in humans. Thus, we sought to continuously assess bone destruction using radiography in tumor-bearing mice treated with vehicle or DMXAA (20 mg/kg, i.p. at d3 and d7). The grade of bone destruction was scored on a range from 1 to 5 using high-resolution X-ray radiographs of tumor bearing femora, as described by Honore et al ^33^. DMXAA treatment decreased the bone destruction score on d8, d11 and d15 after LLC inoculation compared with vehicle group (**Fig. 2a-b**). No sex differences were observed in the protective effects of DMXAA on bone destruction (**Extended Data Fig. 1f**). To explore the microarchitecture of bone, we also employed micro computed tomography (Micro-CT) with 3-dimentional reconstruction analysis *ex vivo* on the distal aspect of tumor-bearing femurs. On d11 after tumor implantation and vehicle or DMXAA treatment as in **Fig. 2a**, 3D reconstruction showed less bone cancer-induced trabecular bone loss and fewer cortical bone lesions in DMXAA-treated mice compared with vehicle group (**Fig. 2c**). Quantitative assessments for bone microstructural parameters demonstrate there is higher trabecular bone connectivity density (Conn.D) and increased cortical bone volume/total volume (BV/TV) in mice administered DMXAA (**Fig. 2d**). Next, we analyzed bone destruction by X-ray radiography following treatment with ADU-S100 or ZA (d3 and d10, as in **Fig. 1f**). Similar to DMXAA, both ADU-S100 and ZA treatment reduced the bone destruction score at d11 and d15 after tumor inoculation (**Fig. 2e-f**).

**Figure 2.**
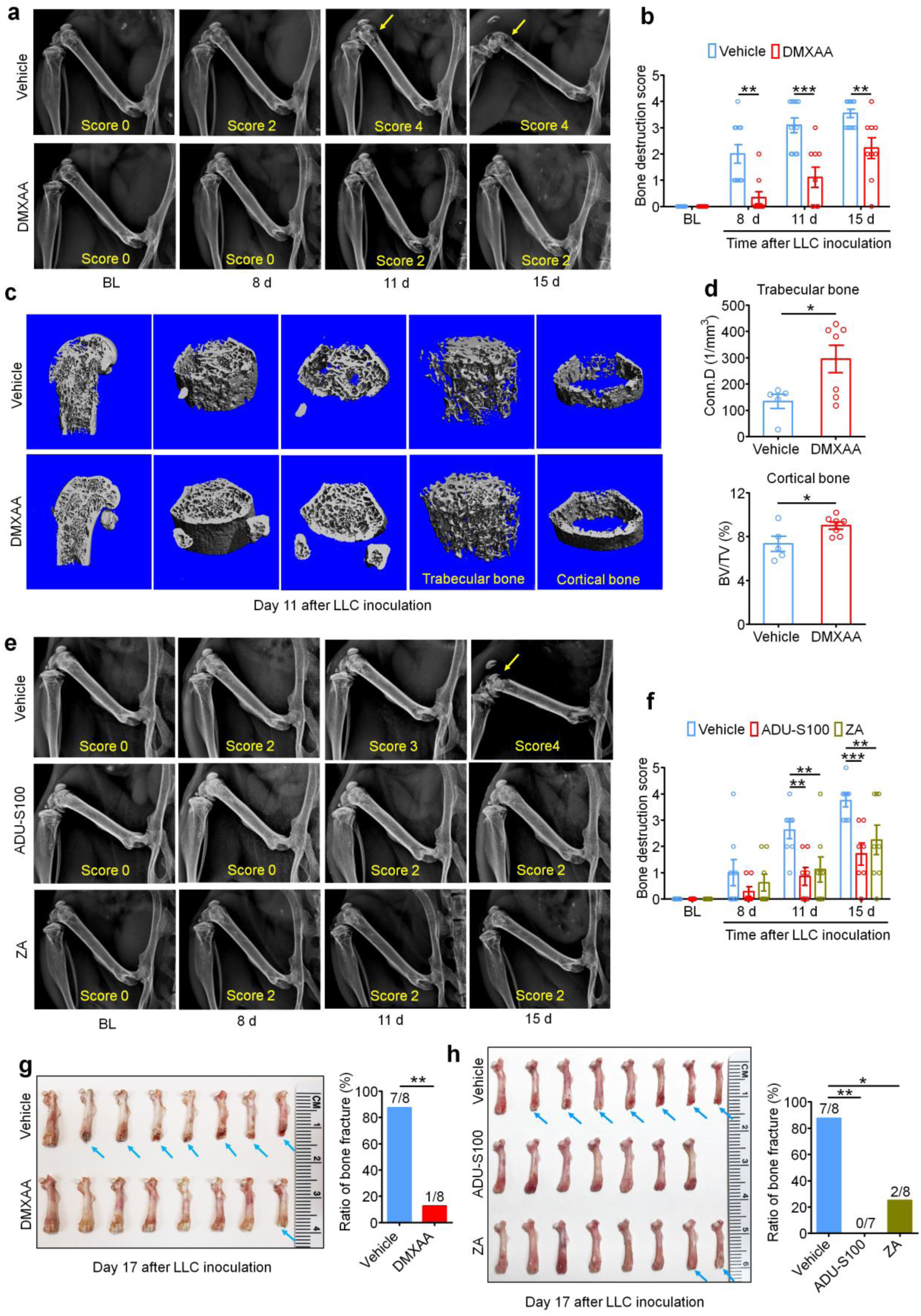
STING agonists attenuate cancer-induced bone destruction. **a.** Representative radiographs of tumor-bearing femora from vehicle or DMXAA treated mice. Bone destruction score is indicated in each image and arrows show bone lesions with scores over 3. **b.** Quantification for (**a**) (n = 9-11 mice/group). **c.** Micro-CT images showing trabecular and cortical bone destruction in the distal part of tumor-bearing femora on d11 after LLC inoculation. **d.** Morphometric quantification of micro-CT images with analysis of trabecular bone (Conn.D; upper) or cortical bone (BV/TV) in LLC inoculated femora from vehicle or DMXAA-treated mice (n = 5-7 mice/group). **e-f.** Radiographical analysis of bone destruction in mice administered vehicle, ADU-S100, or ZA at the indicated timepoints after tumor inoculation. (**e**) Representative X-ray images. Bone destruction score is labeled on the bottom of each photo and arrow indicates bone destruction score more than 3. (**f**) Quantification of images in panel e (n = 7-8 mice/group). **g.** Images of femurs and quantification of the proportion with bone fracture from tumor bearing femora taken from vehicle or DMXAA-treated mice on d17 after LLC inoculation. Arrows indicate the disconnection and absence of the distal part of the femora (n = 8 mice/group). **h.** Images of tumor bearing femora with indicated treatment harvested on d17 after LLC inoculation (left) and quantification of the proportion with bone fracture (right). Arrows indicate lesion and loss of the distal aspect of the femora (n=7-8 mice/group as indicated). Data indicate the mean ± SEM. \**P* < 0.05, \*\**P* < 0.01 and \*\*\**P* < 0.001, repeated-measures two-way ANOVA with Bonferroni’s *post-hoc* test (**b, f**); two-tailed Student’s t-test (**d**); Fisher’s exact test (**g, h**).

Development and progression of cancer-induced osteolytic bone destruction frequently leads to bone fracture, which is an important component of SREs in patients with bone metastasis and is associated with decreased overall survival ^34^. On d17 after LLC inoculation, mice were euthanized and the tumor bearing femora were collected and the distal tumor-bearing femur where bone destruction occurs was analyzed. Notably, we found that 87.5% (7/8 mice) of vehicle-treated mice suffered bone fractures, whereas only 12.5% (1/8 mice) mice treated with DMXAA developed distal bone fractures (**Fig. 2g**). Likewise, 0% (0/7 mice) in the ADU-S100-treated group and 25% (2/8 mice) in the ZA group developed bone fractures (**Fig. 2h**), indicating both STING agonists and ZA could significantly reduce bone destruction.

### STING agonist treatment protects against breast cancer induced bone pain and bone destruction

Similarly to lung cancer, breast cancer is also prone to metastasize to bones and cause bone destruction ^7^. To explore the potential protective effect of STING agonists in breast cancer-induced bone destruction, we utilized the E0771 medullary breast carcinoma cell line to establish a syngeneic mouse model of breast cancer-induced bone cancer pain in female C57BL/6 mice. Similar to the LLC line, tumors established with the E0771 line also induce osteolytic bone lesions ^35^. After intra-femur inoculation, mice were treated with vehicle, DMXAA or ADU-S100 followed by behavioral testing and X-ray radiography of tumor-bearing femurs (**Fig. 3a**). Similar to our results in the LLC bone cancer pain model, we found that DMXAA and ADU-S100 treatment could markedly reduce mechanical allodynia, cold allodynia and spontaneous pain compared to vehicle treatment (**Fig. 3b-d**) but had no effect on body weight (**Fig. 3e**). Furthermore, both DMXAA and ADU-S100 could also attenuate bone destruction scored from X-ray images of the E0771-bearing femora (**Fig. 3f-g**). Thus, STING agonists can protect against cancer-induced bone pain and bone destruction caused by multiple cancer subtypes prone to bone metastasis.

**Figure 3.**
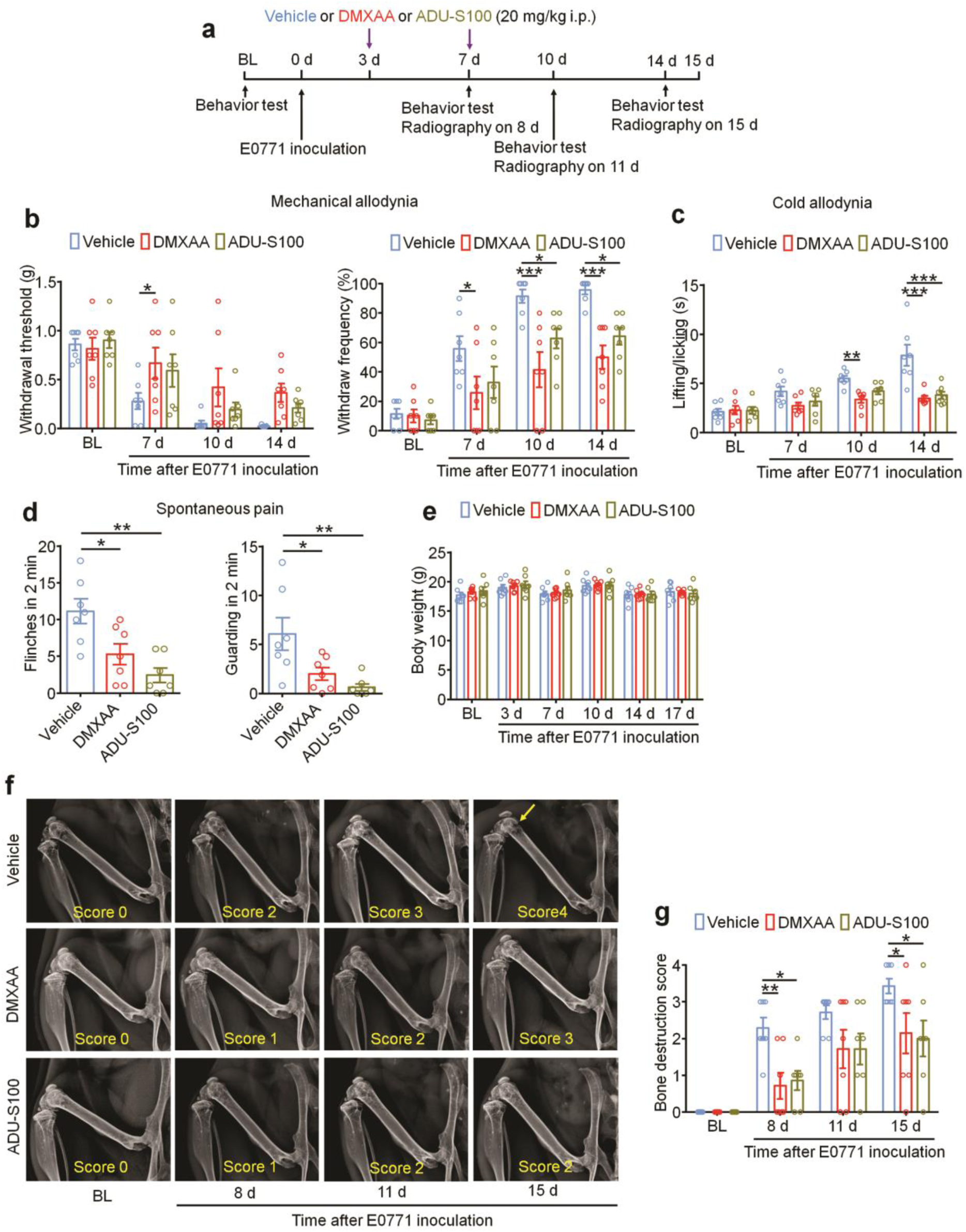
STING agonists confer protection in bone cancer induced by the breast cancer cell line E0771. **a.** Experimental diagram indicating vehicle, DMXAA, or ADU-S100 treatment, behavioral testing, and radiography. **b.** von Frey testing to determine withdrawal threshold (left) and frequency (right) from female mice treated with vehicle, DMXAA or ADU-S100 (n = 7 mice/group). **c.** Cold allodynia testing performed at BL, d7, d10 and d14 after E0771 inoculation (n = 7 mice/group). **d.** Spontaneous pain as quantified by number of flinches (left) or guarding (right) behaviors over a 2 minute interval at d14 after E0771 inoculation in mice with the indicated treatments (n = 7 mice/group). **e.** Measurement of body weight in mice with the indicated treatment (n = 7 mice/group). **f-g.** Representative X-ray images of tumor bearing femora (**f**) and quantification (**g**) of bone destruction score from radiography test on BL, d8, d11 and d15 post-E0771 implantation (n = 7 mice/group). Bone destruction score is indicated in each image and arrows show bone lesions with scores over 3. Data indicate the Mean ± SEM. \**P* < 0.05, \*\**P* < 0.01 and \*\*\**P* < 0.001, repeated-measures two-way ANOVA with Bonferroni’s *post hoc* test (**b, c, e, g**); one-way ANOVA with Bonferroni’s *post-hoc* test (**d**).

### Protective effect of DMXAA on bone pain and bone destruction is STING dependent

To verify the antinociceptive and bone anabolic effects of DMXAA are mediated by STING, WT mice and STING “goldenticket” knockout (STING*^gt/gt^*) mice ^36^ were inoculated with LLC cells intrafemorally followed by vehicle or DMXAA (20 mg/kg) administration (i.p.) on d3 and d7 post LLC injection. Notably, STING*^gt/gt^* mice displayed markedly reduced hindpaw withdrawal threshold and increased withdrawal frequency in von Frey tests compared to WT mice at baseline. DMXAA treatment significantly attenuated mechanical and cold allodynia in WT mice, and this effect was abolished in STING*^gt/gt^* mice (**Extended Data Fig. 2a-b**). We also measured cancer-induced bone destruction in these mice using radiographic examination of bone destruction of the tumor-bearing distal femurs. We observed a reduction in the bone destruction score in DMXAA-treated WT mice at d11 and d15 after tumor inoculation, and this effect was abolished in STING*^gt/gt^* mice (**Extended Data Fig. 2c-d**). We did not see body weight changes after the experimental manipulations (**Extended Data Fig. 2e)**. Thus, as expected, the protective effects of DMXAA on cancer-induced pain and bone destruction are mediated by STING.

### IFN-I signaling mediates the protective effects of STING agonists in bone cancer

STING activation leads to the transcriptional induction of interferon response genes and the robust production and release of type-I interferons, including IFN-α and IFN-β. To confirm that systemic administration of STING agonists leads to increased IFN-I response both systemically and locally within the tumor microenvironment, we analyzed the level of IFN-α and IFN-β by ELISA. We found that serum levels of IFN-α increased approximately 1000-fold 4h after a single i.p. injection of DMXAA (20 mg/kg) or ADU-S100 (20 mg/kg) on d3 after tumor inoculation compared to vehicle group, and this increase was maintained for up to 24 hours. Meanwhile, serum IFN-β levels were also dramatically upregulated 4h after DMXAA and ADU-S100 administration (**Fig. 4a**). On d3 after LLC implantation, the bone marrow (BM) from tumor bearing femora were also collected 4h after vehicle, DMXAA or ADU-S100 i.p. treatment and analyzed by ELISA. Both IFN-α and IFN-β were sharply increased in BM lysate in mice treated with DMXAA or ADU-S100 (**Fig. 4b**). Thus, systemic administration of STING agonists promoted a robust IFN-I response systemically and in the bone cancer tumor microenvironment.

**Figure 4.**
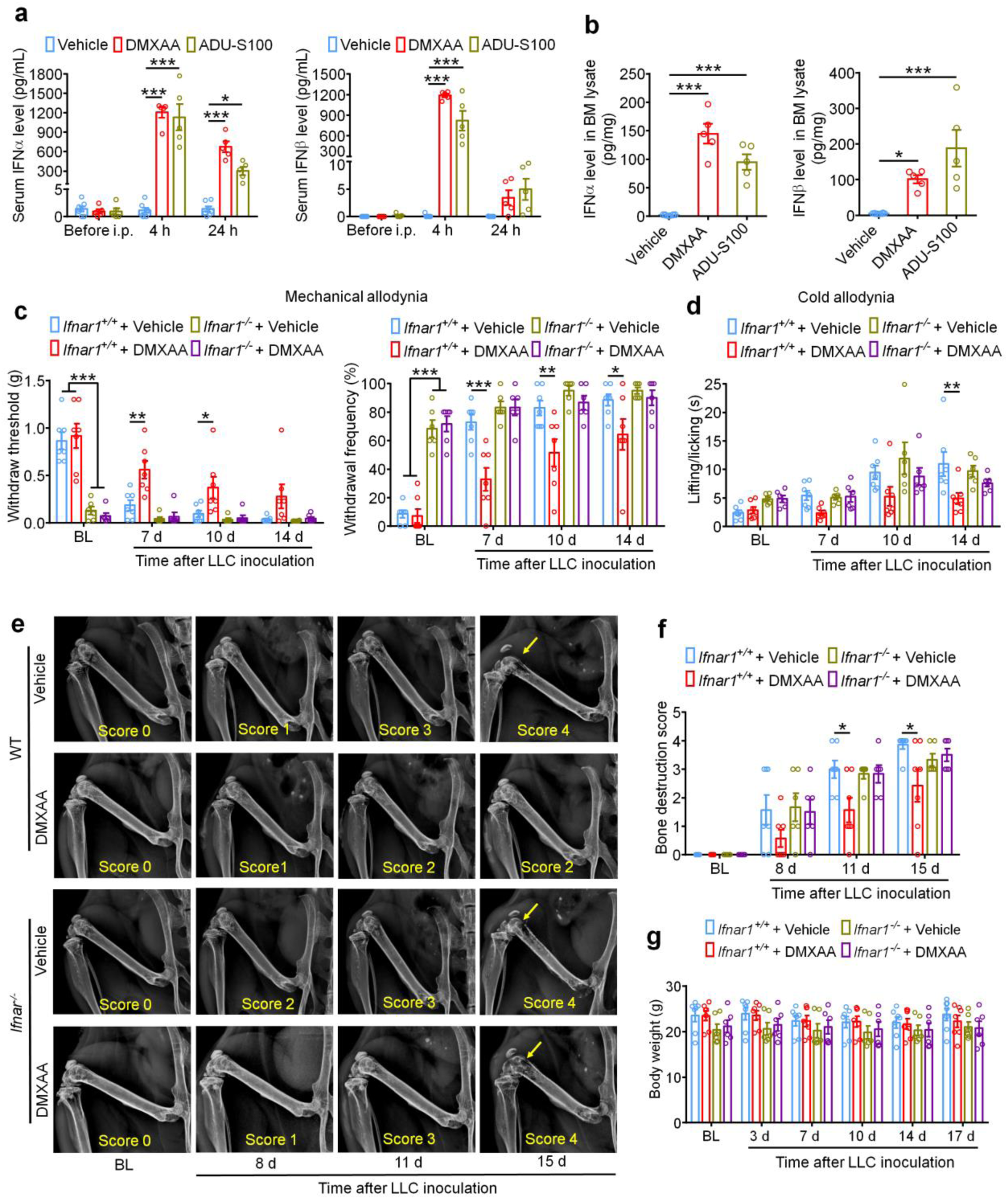
The protective effects of STING agonists are mediated by host IFN-I signaling. **a.** Analysis of serum IFN-α (left) and IFN-β (right) levels at BL and again 4h or 24h after DMXAA or ADU-S100 treatment on d3 after tumor implantation (n = 5 mice/group). **b.** IFN-α (left) and IFN-β (right) levels in bone marrow (BM) lysates from mice treated with vehicle, DMXAA or ADU-S100 (n = 5). **c.** von Frey testing to determine withdrawal threshold (left) and frequency (right) from *Ifnar1^+/+^* or *Ifnar1^−/−^* mice treated with vehicle or DMXAA (2 x 20mg/kg, i.p.; n = 6-7 mice/group). **d.** Analysis of cold allodynia in vehicle or DMXAA-treated *Ifnar1^+/+^* mice and *Ifnar1^−/−^* mice (n = 6-7 mice/group). **e-f.** Bone destruction scores from radiographs of tumor bearing femora in *Ifnar1^+/+^* and *Ifnar1^-/-^* mice with the indicated treatment on d0, d8, d11 and d15 after LLC injection. Arrows show bone lesions with destruction scores over 3. (**e**) Representative X-ray images. (**f**) Quantification for panel e (n = 6-7 mice/group). (**g**) Body weight measurement after vehicle or DMXAA treatment (n = 6-7 mice/group). Data indicate the mean ± SEM. \**P* < 0.05, \*\**P* < 0.01 and \*\*\**P* < 0.001, repeated-measures two-way ANOVA with Bonferroni’s *post-hoc* test.

The IFN-α/β receptor (IFNAR) is a heterodimeric signal transducing receptor complex composed of Ifnar1 and Ifnar2, each of which is required for IFN-I signaling. To test how IFN-I signaling contributes to the protective effects of STING agonists in the bone cancer model, we again introduced LLC cells into the femora of *Ifnar1^+/+^* (WT) or *Ifnar1^−/−^* (KO) mice to establish the bone cancer models in mice with deficient host IFN-I signaling. Similar to mice lacking STING, we found that *Ifnar1^−/−^* mice exhibited mechanical hypersensitivity at baseline compared to WT littermate control mice (**Fig. 4c**). Following treatment with vehicle or DMXAA (20mg/kg, i.p. at d3 and d7), we found that DMXAA treatment effectively attenuated cancer-induced mechanical allodynia and cold allodynia in WT mice but not *Ifnar1^-/-^* mice (**Fig. 4c-d**). On d11 and d15 after tumor inoculation, DMXAA treatment also significantly reduced the bone destruction score without changing overall body weight in WT mice, but this effect was abolished in *Ifnar1^-/-^* mice (**Fig. 4e-g**). Therefore, host IFN-I signaling through Ifnar1 is required for the protective effects of STING agonists on cancer pain and bone destruction induced by bone cancer.

### DMXAA inhibits bone cancer-induced hyperexcitability of DRG nociceptive neurons

Given that cancer-evoked pain in our bone cancer model is transduced by peripheral nociceptors in the dorsal root ganglion (DRG), we next sought to determine whether the antinociceptive effects of STING agonists in bone cancer pain are due to direct effects on nociceptor excitability. To this end, WT mice were inoculated with LLC cells to establish bone cancer models and lumbar L3-L5 DRGs were isolated on d11 and incubated *ex vivo* with vehicle or DMXAA (30 µM) for 2h followed by patch clamp recordings to measure nociceptor excitability (**Fig. 5a**). Importantly, compared to vehicle, DMXAA incubation of DRGs markedly increased the rheobase of nociceptors, a measure of the current required to evoke action potentials (**Fig. 5b-c**). In addition, we found that bone cancer increased neuronal excitability and acute DMXAA incubation sharply reduced the cancer-induced increase in current-evoked action potential firing compared with vehicle-treated DRGs (**Fig. 5d-e**). Taken together, these data indicate that STING activation with DMXAA can suppress cancer-induced hyperexcitability of DRG nociceptors, and cells present within the DRG are sufficient to mediate these effects.

**Figure 5.**
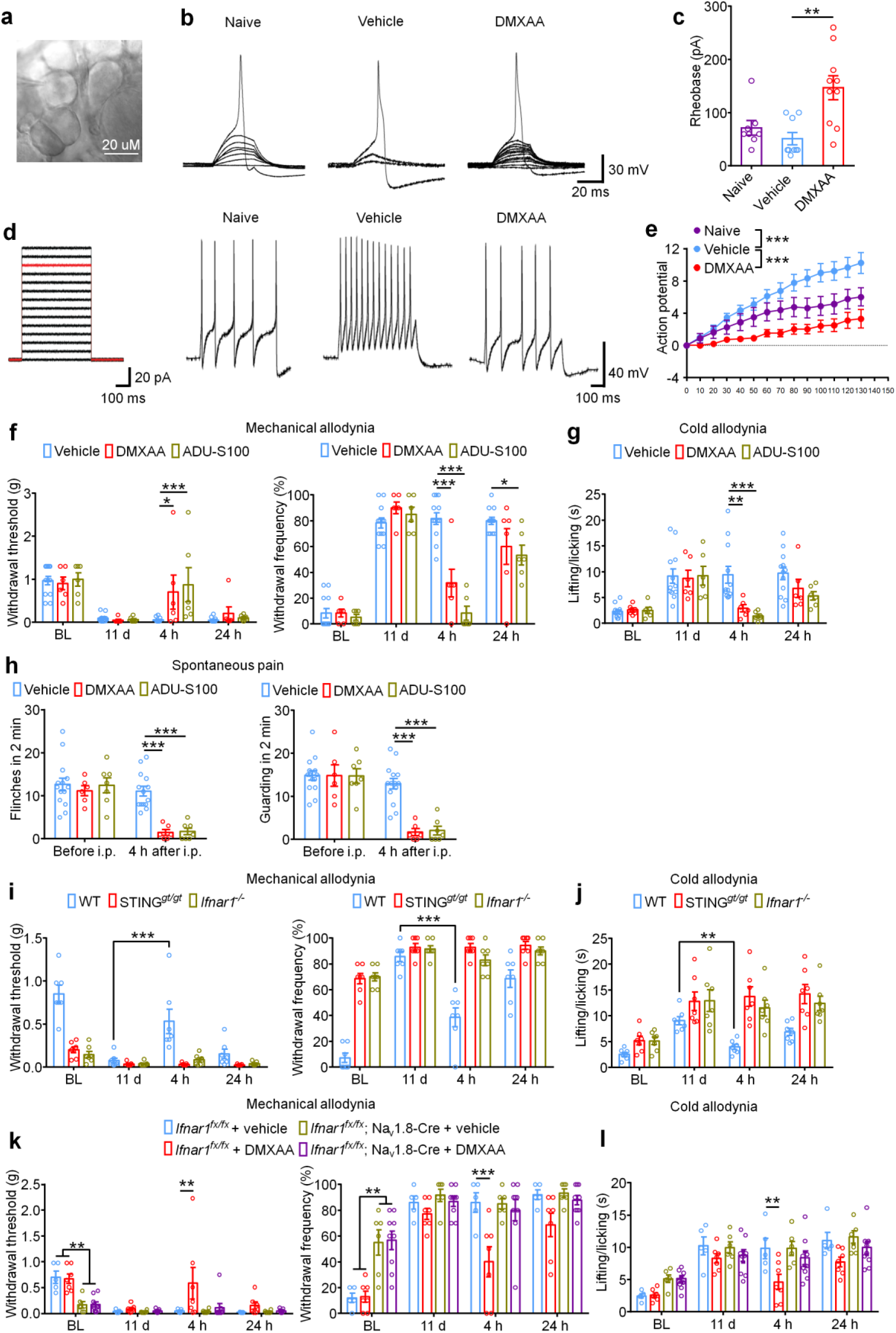
*Ex vivo* STING activation inhibits neuronal hyperexcitability of DRGs from cancer-bearing mice. **a.** Bright field image of a whole-mount DRG shows a recording micropipette sealed on a small-diameter DRG neuron (nociceptor). **b.** Representative traces of rheobases: current clamp recordings of the membrane potential from small diameter DRG neurons of naïve mice or bone cancer-bearing animals with or without DMXAA (30 µM, 2h). Current injection for action potential induction starts from 0 pA and increases 10 pA per step for 30 ms. **c.** The averages of rheobases from naïve mice, vehicle or DMXAA-treated group (naïve: n = 8 neurons/4 mice; vehicle: n= 9 neurons/4 mice; DMXAA: n=10 neurons/4 mice). **d.** Left: injected current steps for the induction of action potentials starts from 0 pA and increases 10 pA per step for 300 ms. Right: representative traces showing the response to a 110 pA current injection (red line in left) from small diameter DRG neurons of naïve mice or bone cancer mice with or without DMXAA treatment. **e.** Action potentials in response to increasing current amplitude from naïve mice or bone cancer mice with or without DMXAA (vehicle: n=5 neurons/3 mice; DMXAA: n=8 neurons/4 mice). **f-g.** Measurement of mechanical allodynia as indicated by paw withdrawal threshold (**f**, left) or withdrawal frequency (**f**, right) or cold allodynia (**g**) 4h or 24h after a single i.p. injection of vehicle, DMXAA (20 mg/kg), or ADU-S100 (20 mg/kg) on d11 after LLC inoculation (n = 6-12 mice/group). **h.** Quantification of spontaneous pain behaviors, as indicated from flinching (left) or guarding (right) 4h after a single i.p. injection of DMXAA or ADU-S100 at d11 post-LLC (n = 6- 14 mice/group). **i-j.** Measurement of mechanical allodynia by von Frey testing (**i**) and cold allodynia from acetone response (**j**) after DMXAA i.p. injection in WT, STING*^gt/gt^* or *Ifnar1^−/−^* mice (n=7 mice/group). **k-l.** Measurement of mechanical allodynia by von Frey testing (**k**) and cold allodynia by acetone response duration (**l**) after DMXAA i.p. injection in *Ifnar1^fx/fx^*; Na_v_1.8-Cre (*Ifnar1*-cKO) mice or *Ifnar1^fx/fx^* (WT) mice (n=5-9 mice/group). Data indicate the mean ± SEM. \**P* < 0.05, \*\**P* < 0.01 and \*\*\**P* < 0.001, two-tailed Student’s t-test (c, e); repeated-measures two-way ANOVA with Bonferroni’s *post-hoc* test (f, g, h, i, j, k, l).

Repeated administration of STING agonists may suppress cancer-induced pain by reducing tumor burden, reducing bone destruction, by a neuronal mechanism involving direct suppression of nociceptor activity, or a combination of all three of these mechanisms. Our electrophysiological data indicate that STING agonists can suppress bone cancer-induced pain via a direct neuronal mechanism which is independent to effects on tumor growth or bone destruction. To test whether acute administration of STING agonists can suppress bone cancer-induced pain, we performed behavior tests in mice on d11 after tumor inoculation 4h after single i.p. injection of vehicle, DMXAA, or ADU-S100. Notably, both STING agonists induced a substantial reduction in mechanical allodynia, cold allodynia, and spontaneous pain (**Fig. 5f-h**). Since the natural activators of STING are intracellular double-stranded DNA (dsDNA) and the bacterial cyclic dinucleotide 3’3’-cGAMP^37^, we also assessed whether dsDNA and 3’3’-cGAMP could produce antinociception in the bone cancer model. Interestingly, i.p. administration of dsDNA (30 µg, complexed with LyoVec to facilitate cellular penetration) or cGAMP (20 mg/kg) could attenuate cold allodynia or/and mechanical allodynia 4h after injection on d11 post LLC implantation (**Extended Data Fig. 3a-d**). Additionally, we tested whether the acute antinociceptive effects of DMXAA were STING- and Ifnar1-dependent by injecting DMXAA (20 mg/kg, i.p.) into WT, STING*^gt/gt^* mice and *Ifnar1^−/−^* mice, measuring mechanical and cold allodynia 4h after injection. We found DMXAA could reduce mechanical allodynia and cold allodynia in WT mice but not in STING*^gt/gt^* mice or *Ifnar1^−/−^* mice (**Fig. 5i-j**). To further determine whether neuronal IFN-I signaling is responsible for the acute antinociceptive effects of STING agonists, we established the bone cancer pain model using mice lacking *Ifnar1* selectively in sensory neurons (*Ifnar1^fx/fx^*; Na_v_1.8-Cre; *Ifnar1*-cKO), or their wildtype littermates. Importantly, we found the antinociceptive effects conferred by a single administration of DMXAA (20 mg/kg, i.p.) were present in WT, but not *Ifnar1*-cKO littermates (**Fig. 5k-l**). Given the immediacy of these effects, and taken in conjunction with our electrophysiological data, we conclude that STING agonists exert antinociceptive effects via direct actions on nociceptors in an Ifnar1-dependent mechanism.

### STING agonists suppress local bone cancer tumor burden and further metastasis

Intratumor injection of STING agonists have been reported to reduce tumor growth by promoting T cell-mediated antitumor immunity in several preclinical animal studies ^11, 19, 38, 39^. It is unknown, however, whether systemic administration of STING agonists can attenuate tumor progression in the bone marrow, which is generally regarded as an overwhelmingly immunosuppressive tumor microenvironment ^21^. To answer this question, luciferase-labeled LLC cells (LL/2-LUC2 cell line) were used to establish the metastatic bone cancer model via intrafemoral inoculation, thereby enabling measurement of local tumor burden by *in vivo* bioluminescent imaging. Mice were treated with vehicle or DMXAA (20 mg/kg, i.p. at d3 and d7), followed by *in vivo* bioluminescence imaging at d8, d11, and d15. Notably, mice treated with DMXAA exhibited lower local tumor burden at d11 and d15, as measured by total flux of LL/2-Luc2 cells in tumor-bearing mice (**Fig. 6a**). By d17, tumor growth beyond the normal anatomic boundaries of the distal femur could be visually observed, leading to an increase in the circumference of the tumor-inoculated (ipsilateral) thigh compared to the contralateral side. To quantify this, we measured the ratio of the maximum thigh circumference (ipsilateral/contralateral), which accurately reflects local tumor volume ^40^. Notably, we found that DMXAA and ADU-S100 treatment, but not ZA treatment, could reduce the ratio of maximum thigh circumference compared to the vehicle-treated group on day 17 in both the LLC and E0771-induced bone cancer models (**Fig. 6b-c**). To test whether these effects were dependent on host-intrinsic STING and Ifnar1, we measured local tumor burden using thigh circumference at d17 in STING*^gt/gt^* and *Ifnar1^−/−^* mice and found that this protective effect was abolished in mice lacking either STING or *Ifnar1* (**Extended Data Fig. 4a-b**). Thus, systemic administration of STING agonists reduces local bone cancer tumor burden in a STING- and Ifnar1- dependent manner.

**Figure 6.**
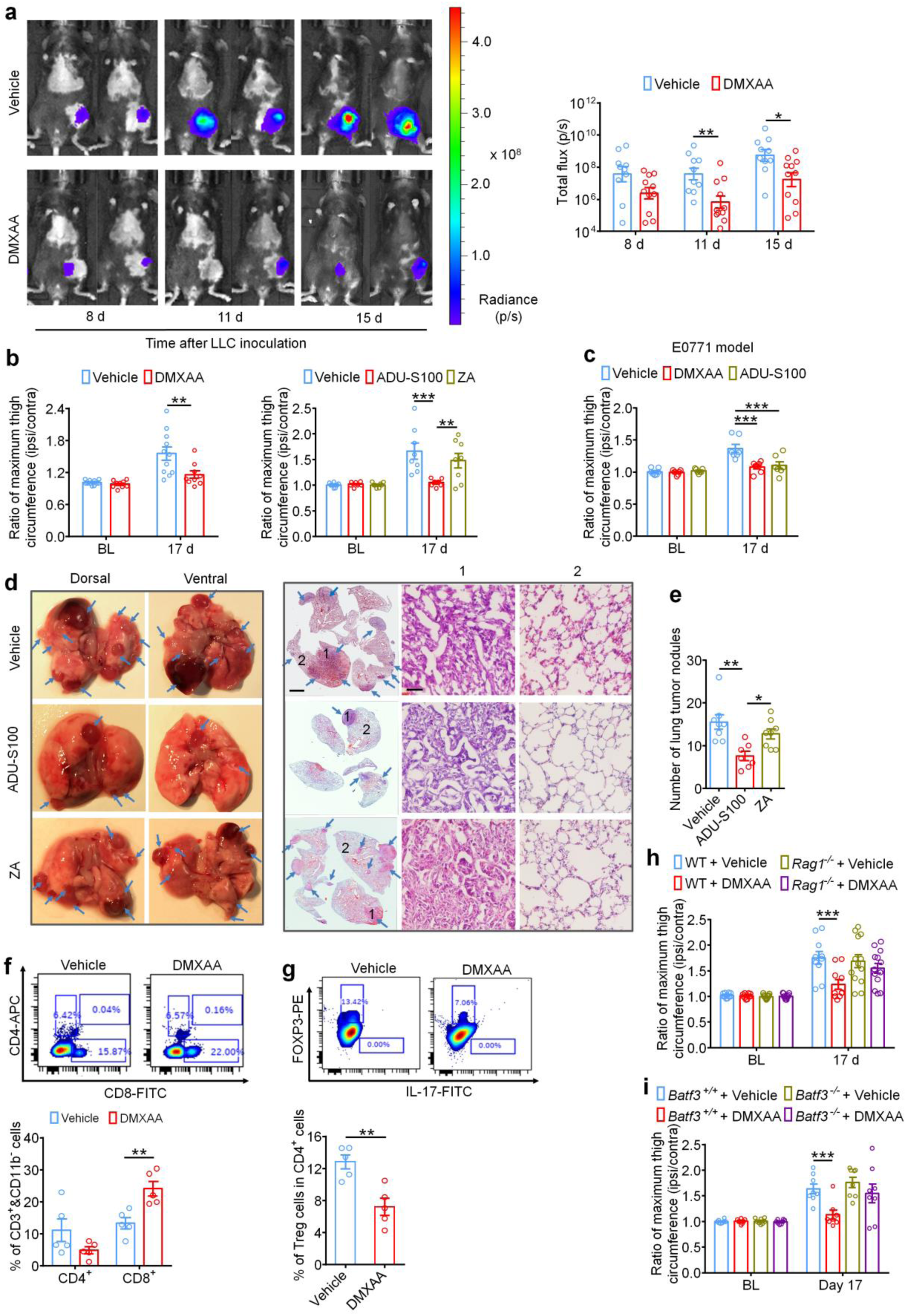
Systemic STING agonists reduce local tumor burden in the bone cancer tumor microenvironment. **a.** *In vivo* bioluminescence of flux emitted by LL/2-Luc2 cells in tumor bearing femur after vehicle or DMXAA treatment (2 x 20 mg/kg, i.p.) measured at d8, d11, and d15 post tumor inoculation (n = 10-11 mice/group). Images (left) were obtained at 15 min after i.p. injection of D-luciferin (30 mg/kg). Right, quantification of a. **b.** Ratio of maximum thigh circumference reflecting local tumor burden in mice with each indicated treatment on d17 after LLC implantation. Left, vehicle or DMXAA treatment (2 x 20 mg/kg, i.p.; n = 9-11 mice/group). Right, vehicle, ADU-S100 (2 x 20 mg/kg, i.p.) or ZA (2 x 100 µg/kg, i.p.) treatment (n = 7-8 mice/group). **c.** Ratio of maximum thigh circumference in mice administered with vehicle, DMXAA or ADU-S100 (2 x 100 µg/kg, i.p.) on d17 after implantation of E0771 breast cancer cells (n = 7 mice/group). **d.** Images of lung tumor nodules in mice with each indicated treatment on d17 after LLC inoculation. Left, representative dorsal and ventral murine lung image, with arrows showing metastatic tumor nodules. Right, H&E staining for sections from lung samples in Left. Arrows indicates the areas with tumor cells, scale bar, 2 mm. 1 and 2 are enlarged images showing tumor tissue and peritumoral areas, respectively. Scale bar, 50 µm. Note that tumor cells have large and irregular nuclei with loss of the normal alveolar structure. **e.** Quantification of panel d (n = 7-8 mice/group). **f-g.** FACS analysis of CD4^+^ and CD8^+^ T cells (**f**) or Treg cells (**g**) within the bone marrow tumor microenvironment in mice treated with vehicle or DMXAA (2 x 20 mg/kg, i.p.) on d8 post-LLC inoculation (n = 5 mice/group). **h-i.** Local tumor burden as determined by the ratio of maximum thigh circumference in WT and *Rag1^−/−^* (**h**) or *Batf3^+/+^* and *Batf3^−/−^* (**i**) mice with indicated treatment measured on d17 after LLC implantation. Data indicate the mean ± SEM. \**P* < 0.05, \*\**P* < 0.01 and \*\*\**P* < 0.001, repeated-measures two-way ANOVA with Bonferroni’s *post hoc* test (**a, b, c, h, i**); one-way ANOVA with Bonferroni’s *post hoc* test (**e**); two-tailed Student’s t-test (**f, g**).

LLC is a murine lung adenocarcinoma cell line which has affinity to metastasize from the original injection site to pulmonary lobes and form visible tumor nodules ^41^, enabling use of this phenomenon as a measure of metastasis in our model. To test whether systemic administration of ADU-S100 (20 mg/kg i.p.) or ZA (100 µg/kg i.p.) at d3 and d7 could reduce lung metastasis, we analyzed lungs from vehicle-, ADU-S100-, or ZA-treated mice at d17 after intrafemoral LLC inoculation. We found that mice receiving ADU-S100 exhibited fewer lung tumor nodules compared to mice treated with vehicle or ZA (**Fig. 6d-e**). Thus, systemic STING activation with ADU-S100 can inhibit both local tumor burden as well as further tumor metastasis.

Mechanistically, the antitumor effects of STING pathway activation are chiefly attributed to antigen presenting cell (APC)-mediated activation of CD8^+^ T cells ^42^. To test whether systemic STING activation can promote CD8^+^ T cell infiltration into the immunosuppressive tumor microenvironment of the bone marrow in our bone cancer model, mice were administered DMXAA (20 mg/kg i.p.) at d3 and d7 and bone marrow was collected from tumor-bearing femora 24 h after the second DMXAA injection for analysis of tumor-infiltrating lymphocytes (TILs) by flow cytometry. Importantly, we found that DMXAA treatment significantly increased the proportion of (CD11b^-^ CD3^+^) CD8^+^ T cells without significantly changing the proportion of (CD11b^-^ CD3^+^) CD4^+^ T cells in the bone marrow tumor microenvironment (**Fig. 6f**). We further analyzed the proportion of immunosuppressive (CD3^+^ CD4^+^) Foxp3^+^, IL-17^-^ T^reg^ cells and found that DMXAA treatment decreased the proportion of T^reg^ cells in the bone marrow (**Fig. 6g)**. To test whether STING agonist-induced reduction in tumor burden is due to T cell-mediated antitumor immunity, we introduced LLC cells into the intrafemoral cavity of WT or *Rag1^-/-^* mice lacking mature B and T cells, followed by vehicle or DMXAA treatment (20 mg/kg i.p. at d3 and d7 post-inoculation) and measurement of maximum thigh circumference at d17 as in **Fig. 6b**. We found that DMXAA effectively reduced the ratio of maximum thigh circumference only in WT mice but not in *Rag1^−/−^* mice (**Fig. 6h**). Next, given that conventional type 1 dendritic cells (cDC1) have been demonstrated to be critical for cross-priming adaptive T cell responses against tumors through STING-mediated IFN-I induction, we also utilized cDC1-deficient *Batf3^−/−^* mice to test whether the acute and/or long-term protective effects would depend on adaptive antitumor immunity. As expected, upon femoral inoculation of *Batf3^+/+^* or *Batf3^-/-^* mice with LLC cells, only tumor-bearing *Batf3^+/+^* mice but not *Batf3^−/−^* mice exhibited a decrease in local tumor burden at d17 following DMXAA treatment (**Fig. 6i**). Thus, we conclude that systemic activation of host-intrinsic STING-mediated IFN-I signaling facilitates antitumor immunity by promoting TIL entry into the normally immunosuppressive bone marrow TME.

### STING agonists inhibit cancer-induced osteoclast differentiation via IFN-I signaling

IFN-α and IFN-β were previously reported to inhibit the differentiation of murine and human preosteoclasts into osteoclasts ^23^. Given our data indicating that STING agonists can reduce bone destruction, we sought to determine whether the bone protective effects are mediated by direct effects on osteoclastogenesis. To this end, we measured osteoclast cell numbers in the distal tumor-bearing femora at d11 after inoculation in mice treated with vehicle or DMXAA (20 mg/kg i.p. at d3 and d7). Notably, DMXAA-treated mice exhibited far significantly fewer osteoclasts (**Fig. 7a**), but no changes were observed in bone-forming osteoblasts (**Fig. 7b**). To further evaluate the activity of osteoclasts and osteoblasts, we collected serum from tumor-bearing mice on BL and d17 after LLC inoculation and measured serum CTX-I and PINP levels, which are markers for bone resorption and bone formation, respectively ^27, 43^. DMXAA could effectively reduce CTX-I levels on d17 but had no effect on serum PINP levels (**Fig. 7c**). These data indicate that STING activation with DMXAA can suppress bone cancer-driven osteoclast formation and their bone catabolic activity.

**Figure 7.**
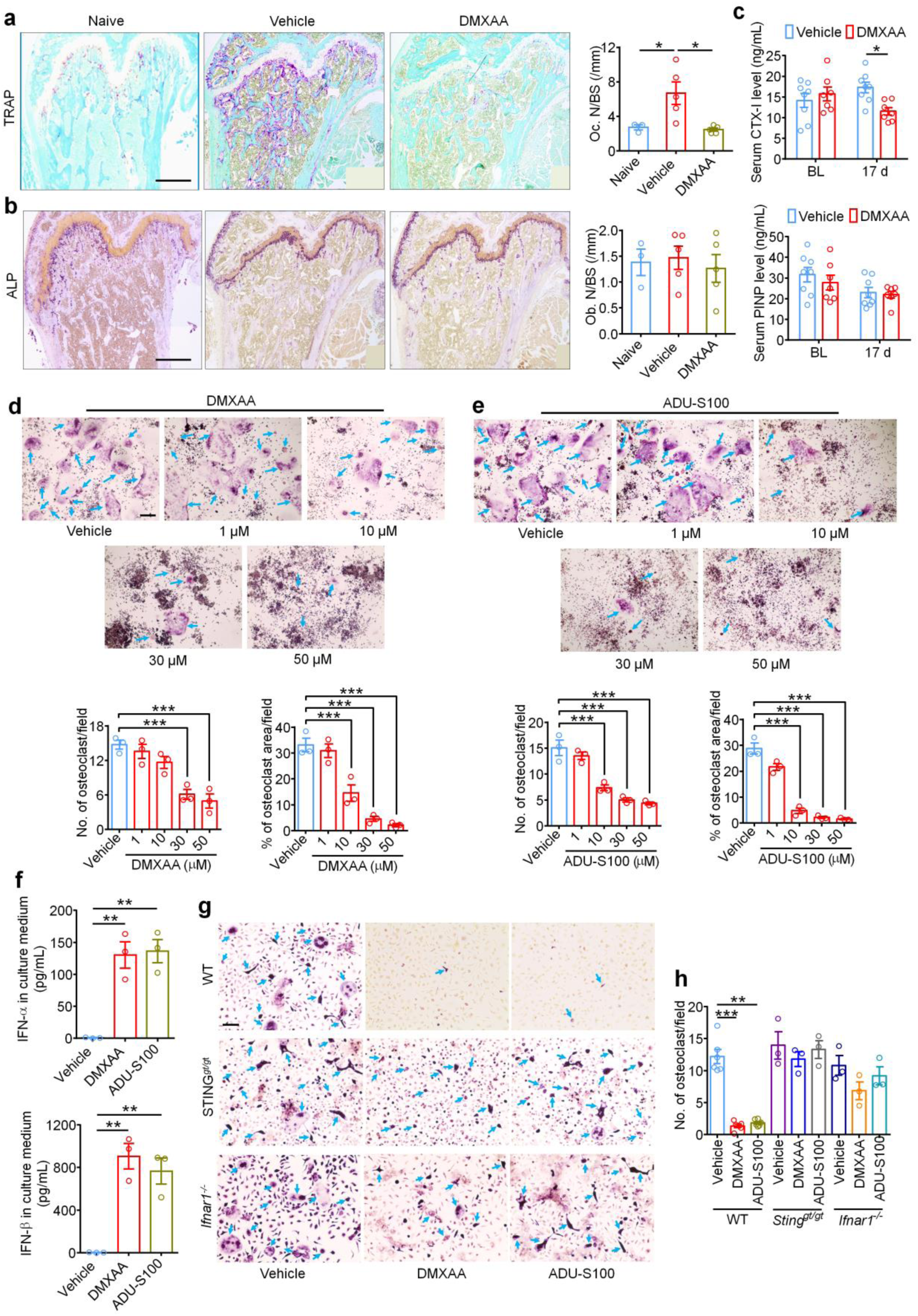
STING agonists attenuate cancer-induced osteoclastogenesis. **a.** Representative images (left) and quantification (right) of TRAP staining to reveal osteoclast numbers in the tumor-bearing distal femora from mice treated with vehicle or DMXAA (2 x 20 mg/kg, i.p.) measured on d11 after LLC inoculation (n = 5 mice/group). Scale bar, 500 µM. **b.** Images (left) and quantification (right) of ALP staining to reveal osteoblasts in the tumor-bearing distal femora at d11 from mice with the indicated treatments (n = 5 mice/group). Scale bar, 500 µM. **c.** Measurement of serum CTX-I and PINP levels by ELISA at BL or d17 in vehicle or DMXAA (2 x 20 mg/kg, i.p.) treated mice (n = 7-8 mice/group). **d-e.** TRAP staining revealing osteoclast numbers after differentiation from RAW264.7 cells stimulated with 35 ng/ml RANKL, in the presence of increasing concentrations of DMXAA (**d**) or ADU-S100 (**e**). Arrows indicate TRAP^+^ multinucleated osteoclasts. Top, representative TRAP stained images. Bottom, quantification (n = 3 independent replicates). Scale bar, 200 µm. **f.** ELISA quantification of IFN-α and IFN-β levels in the culture medium of BMDM cells 24h after DMXAA (30 µM) or ADU- S100 (30 µM) co-incubation. RANKL: 35 ng/ml, MCSF: 20 ng/ml (n = 3 independent experiments). **g-h.** TRAP staining for osteoclasts differentiated from BMDM cells from WT mice, STING*^gt/gt^* mice or *Ifnar1^−/−^* mice, each treated with vehicle, DMXAA (30 µM) or ADU-S100 (30 µM). RANKL: 35 ng/ml, MCSF: 20 ng/ml. (**g**) representative images of TRAP staining. Arrows indicate TRAP^+^ multinucleated osteoclasts. Scale bar, 100 µm. (**h**) quantification for (**g**) (n = 3-6 individual cultures/group). Data indicate the mean ± SEM. \**P* < 0.05, \*\**P* < 0.01 and \*\*\**P* < 0.001, two-tailed Student’s t-test (**a, b**); repeated-measures two-way ANOVA with Bonferroni’s *post-hoc* test (**c**); one-way ANOVA with Bonferroni’s *post hoc* test (**d, e**).

Given that systemic STING agonist treatment reduces osteoclast numbers *in vivo*, we sought to determine whether STING pathway activation can promote osteoclast differentiation *in vitro*. Murine macrophage RAW 264.7 cells were treated with RANKL (35 ng/ml, for 6 days) to promote osteoclast differentiation ^27^ in the presence of vehicle or an escalating dose of DMXAA or ADU-S100. Importantly, we found that both DMXAA and ADU-S100 dose dependently inhibited osteoclast differentiation (**Fig. 7d-e**). Bone marrow cells from WT, STING*^gt/gt^*, or *Ifnar1^-/-^* mice were harvested and differentiated into macrophages with 20 ng/ml M-CSF for 3 days. These bone marrow-derived macrophages (BMDM) were further induced into osteoclasts with 20 ng/ml M-CSF and 35 ng/ml RANKL for 7 days ^27^. We collected the BMDM culture medium 24h after incubation with DMXAA or ADU-S100 and found both agonists induced a drastic increase in IFN-α and IFN-β levels in the culture medium, although IFN-β induction was much greater (**Fig. 7f**). Furthermore, TRAP staining showed that DMXAA or ADU-S100 treatment (30 µM each) could significantly inhibit osteoclast formation from BMDM from WT mice but not from STING*^gt/gt^* or *Ifnar1^-/-^* mice (**Fig. 7g-h**). To provide further support that these effects were reliant on IFN-α/β signaling, anti-IFN-α (600 ng/ml) or anti-IFN-β (600 ng/ml) neutralizing antibodies were added to the induction medium of BMDM followed by analysis of osteoclast formation. We found that anti-IFN-β antibody could block the inhibition of osteoclast formation by DMXAA or ADU-S100 (**Extended Data Fig. 5a-d**). Taken together, these data indicate that activation of the STING/IFN-I signaling axis can reduce osteoclastogenesis.

Our findings indicate that STING agonists produce antinociception, reduce tumor burden, and reduce bone destruction and osteoclastogenesis. One could argue that both the antinociceptive effects and the bone protective effects are secondary to T cell-mediated antitumor immunity. To test this possibility, we again introduced LLC cells into the intrafemoral cavity of WT or *Rag1^-/-^* mice, followed by vehicle or DMXAA treatment (20 mg/kg i.p.) at d3 and d7 post-inoculation. Notably, bone cancer-induced mechanical and cold allodynia were reduced by DMXAA treatment in both WT and *Rag1^-/-^* mice at early stages (7d and 10d), but not at later stages (14d; **Extended Data Fig. 6a-b**). Likewise, DMXAA treatment led to an improvement in the bone destruction score in both WT and *Rag1^-/-^* mice at d11, but not at d15 and at d17 for bone fracture (**Extended Data Fig. 6c-e**). Thus, these data indicate that DMXAA suppresses pain and bone destruction in a T cell-independent mechanism at early stages, and thus, these effects are likely due to direct suppression of nociceptor excitability and osteoclastogenesis. As the local tumor burden increases at later stages, T cell-mediated antitumor immunity may become essential in controlling pain and bone destruction. To further verify these findings, we used *Batf3^+/+^* and *Batf3^−/−^* mice to establish bone cancer pain model. DMXAA treatment (20 mg/kg i.p., d3 and d7) could attenuate mechanical allodynia or cold allodynia in *Batf3^−/−^* mice on d7 and d10 but not day 14 after tumor inoculation (**Extended Data Fig. 7a-b)**. DMXAA also reduced bone destruction on d8 and d11 but not d15 in *Batf3^−/−^* mice (**Extended Data Fig. 7c-d).** These data provide an additional line of evidence that DMXAA suppresses pain and bone destruction in a T cell-independent mechanism at early stages. Overall, we propose a mechanism by which STING agonists induce robust production of type-I interferons, which directly suppress nociceptor excitability and osteoclastogenesis while concurrently promoting T cell-mediated antitumor immunity. Thus, we posit that STING agonists can acutely suppress cancer pain through direct effects, while providing long term relief from bone cancer-induced pain by suppressing osteoclast-mediated bone destruction and relieving local tumor burden (**Fig. 8**).

**Figure 8.**
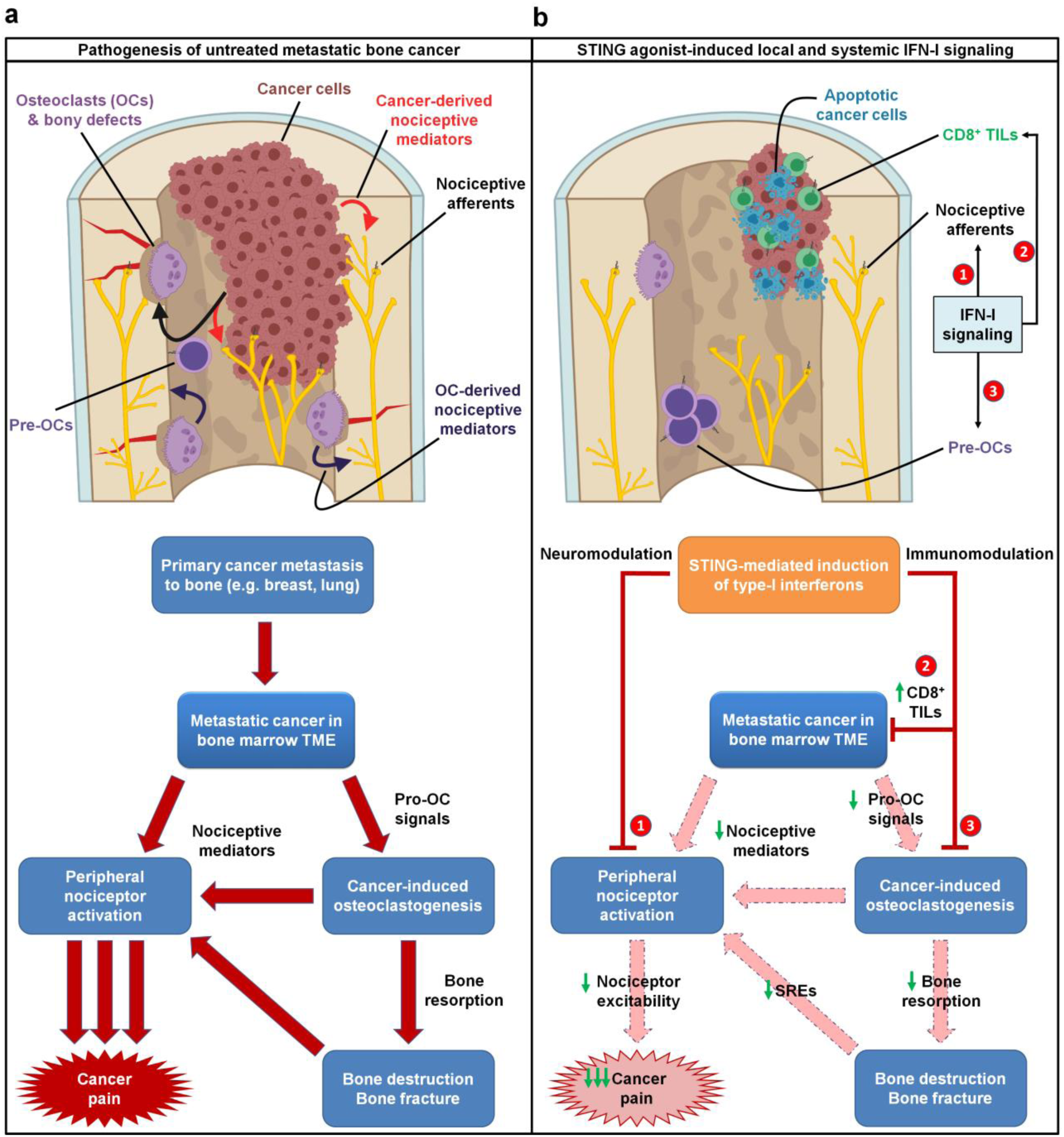
Schematic of mechanisms by which STING agonists directly and indirectly attenuate bone cancer-induced pain. **a.** In untreated metastatic bone cancer, the pathophysiology driving cancer pain is multifaceted. Metastatic cancer cells present within the bone marrow tumor microenvironment (TME) produce pro-osteoclastogenic signals, driving cancer-induced osteoclastogenesis and osteoclast overactivation. Both cancer cells and osteoclasts produce nociceptive mediators, which directly activate peripheral nociceptive afferents present in the TME to produce pain through a direct mechanism. Additionally, cancer-induced osteoclast overactivation also leads to increased bone resorption, leading to increased bone destruction and bony fractures, which also produce pain through nociceptor activation. **b.** STING agonists dramatically attenuate metastatic bone cancer-associated pain through multiple mechanisms, each mediated by host-intrinsic type-I interferon signaling. First, STING-mediated IFN-I signaling directly suppresses excitability of peripheral nociceptors (neuromodulation), leading to acute suppression of pain for the duration of the IFN-I response. In addition, STING-mediated IFN-I signaling promotes CD8^+^ T cell migration into the bone marrow TME, augmenting antitumor immunity and reducing tumor burden. In addition, IFN-I drives suppression of cancer-induced osteoclastogenesis. These two non-neuronal, immunomodulatory effects lead to sustained inhibition of pain by (1) reducing cancer cell- and osteoclast-derived pro-nociceptive mediators and (2) reducing osteoclast-mediated bone resorption, thereby attenuating subsequent bone destruction and bony fractures that frequently evoke pain and skeletal-related events (SRE). Thus, the immunomodulatory and neuromodulatory effects of STING agonists each individually suppress pain through actions on different cell types, and also synergistically suppress cancer pain through a convergence of shared downstream actions.

## Discussion

Our findings elucidate a unique and synergistic advantage of STING agonism as a therapeutic strategy in treating metastatic bone cancer pain and its comorbidities, including tumor-induced bone destruction and functional impairment (e.g., immobility). To our knowledge, this is the first established or prospective immunotherapy agent demonstrated to possess antitumor, antinociceptive, and bone anti-catabolic properties. Notably, while we demonstrate that STING-mediated IFN-I signaling exerts direct effects on pain via direct suppression of nociceptor excitability (neuromodulation), the potent analgesic effects of STING agonists *in vivo* are likely owed to a combinatorial suppression of tumor burden and bone destruction (immune modulation), and direct effects on nociceptive sensory neurons. Likewise, we also demonstrate the STING/IFN- I signaling axis directly suppresses osteoclast differentiation, but the bone protective effects conferred by STING agonists *in vivo* are likely mediated by both direct effects and indirect effects (reduced tumor burden). These interrelated and synergistic protective actions of STING agonists (summarized in **Fig. 8**) would seem to be of particular advantage for the treatment of metastatic bone cancer, which offers unique challenges due to the ongoing presence of systemic disease in conjunction with cancer-associated SREs and severe pain which can themselves increase cancer mortality ^44^.

Notably, the majority of clinical trials exploring the efficacy of STING agonists in human patients to date have focused on intratumoral administration, primarily in advanced stage solid tumors. While systemic administration may increase treatment-related adverse effects (TREs), increased toxicity may be acceptable in patients with metastatic disease and/or poor long-term prognosis, especially if coupled with attenuated cancer pain and improved function. Moreover, while the immune-mediated antitumor effects may be maximized by intratumoral administration of STING agonists, this route of administration may not be sufficient to yield the analgesic or bone anabolic effects we demonstrate here. Notably, in human patients STING agonists exhibit little efficacy as a monotherapy but have shown results when used in conjunction with anti-PD-1 immunotherapy regimens. In these studies, however, results are typically measured in terms of objective response rate (ORR) which only considers tumor size reduction. Notably, anti-PD-1 treatment in the LLC-induced murine bone cancer model alleviated cancer pain but did not change tumor burden^27^. Despite the controversy of STING agonists in controlling tumor growth in pre-clinical and clinical studies, our findings highlight the use of STING-based immunotherapies for treating cancer pain, especially bone cancer pain, due to multiple actions of the STING pathway in immune modulation, neuromodulation, and neuro-immune interactions (**Fig. 8**). Thus, clinical trials testing prospective STING immunotherapy agents should also consider measuring cancer-associated comorbidities, including parameters of functional impairment and pain, as improving the quality of life in cancer patients is equally important as extending the duration of survival.

## Methods

### Reagents

The following reagents were used in this study: DMXAA (Cayman Chemical, 14617), ADU-S100 (Chemietek, CT-ADUS100), 3’3’-cGAMP (Invivogen, tlr-nacga), poly(dA:dT)/LyoVec (Invivogen, tlrl-patc), Zoledronic acid (Cayman Chemical, 14984), mouse RANKL protein (R&D systems, 462-TEC), mouse M-CSF (R&D systems, 416-ML), anti-mouse IFN-α neutralizing antibody (PBL Assay Science, 32100-1), anti-mouse IFN-β neutralizing antibody (PBL Assay Science, 32400-1) and rabbit polyclonal IgG control (Biolegend, CTL-4112).

### Animals

Adult C57BL/6J mice (males and females, 8-10 weeks) were used for all behavioral and biochemical studies. STING “goldenticket” knockout mice (Stock No: 017537), Ifnar1 global knockout mice (Stock No: 028288), Rag1 knockout mice (Stock No: 002216), Ifnar1 conditional knockout mice (*Ifnar1^fx/fx^*; Stock No: 028256) and *Batf3^−/−^* mice (Stock No: 013755) were purchased from the Jackson Laboratory and maintained on a C57BL/6J background. Na_v_1.8-Cre mice, also maintained on a C57BL/6J background, were a gift from Rohini Kuner (University of Heidelberg). Behavioral testing was performed comparing STING*^gt/gt^* and *Ifnar1^-/-^* mice with wildtype littermate controls. These mice were maintained at an AAALAC-approved Duke University facility with two to five mice housed in each cage maintained in a 12h light-dark cycle with *ad libitum* access to food and water. Animals were randomly assigned to different experimental groups. Our previous studies using similar types of behavioral and biochemical analyses ^27, 45, 46^ were used to determine sample size. Males and females were used in a sex and age-matched manner, if not otherwise specified in the figure legends. All mouse procedures were approved by the Institutional Animal Care & Use Committee (IACUC) of Duke University. Animal experiments were conducted in strict accordance with the National Institutes of Health Guide for the Care and Use of Laboratory Animals. The numbers of mice used in different experiments are summarized in **Extended Data Table 1**.

### Cell culture

Murine Lewis lung carcinoma cell line LL/2 (LLC1) (ATCC® CRL-1642), luciferase expressing cell line LL/2-Luc2 (ATCC® CRL-1642-LUC2™) and murine monocyte/macrophage cell line RAW 264.7 (ATCC® TIB-71) were obtained from ATCC. The mouse E0771 breast cancer cell line (94A001) was obtained from CH3 BioSystems. Cells were cultured in high glucose (4.5 g/L) Dulbecco’s modified Eagle medium (Gibco, Thermo Fisher Scientific), supplemented with 10% fetal bovine serum (Gibco, Thermo Fisher Scientific) and 1% antibiotic-antimycotic solution (Sigma-Aldrich). These cells were then cultured in the presence of 5% CO2 at 37 °C. Blasticidin (2 µg/ml, Gibco, Thermo Fisher Scientific) was added into LL/2-Luc2 culture medium and removed 3 days before the inoculation of mice. No testing was performed for mycoplasma contamination.

### Bone cancer pain model

The murine cell lines LLC1, LL/2-Luc2 or E0771 were lightly digested using 0.05% trypsin, followed by centrifugation to remove poorly digested cell clusters. Cells were then resuspended in PBS at a concentration of 1×10^8^ cells/ml. The inoculation was performed as previously described ^27, 47^. Briefly, mice were anesthetized with 4% isoflurane and the left leg was shaved and the skin disinfected with 10% povidone-iodine and 75% ethanol. A superficial incision (0.5-1 cm) was made near the knee joint, exposing the patellar ligament. A new 25-gauge needle was inserted at the site of the intercondylar notch of the left femur into the femoral cavity, which was then replaced with a 10 µL microinjection syringe containing a 2 µL suspension of tumor cells (2 × 10^5^) followed by 2 µL absorbable gelatin sponge solution to seal the injection site. The syringe contents were slowly injected into the femoral cavity over a 2-minute interval. To prevent further leakage of tumor cells outside of the bone cavity, the outer injection site was sealed with silicone adhesive (Kwik-Sil, World Precision Instruments, US). Animals with surgery related movement dysfunction or with outside bone tumor injection were excluded from the study.

### Drug treatment

DMXAA was dissolved in sterile PBS containing 0.75% NaHCO_3_ and ADU-S100 was dissolved in sterile PBS into 20 mg/ml, and they were further diluted 10-fold in sterile PBS prior to injection for *in vivo* experiments. All other reagents were dissolved in sterile saline or PBS. For experiments utilizing rabbit anti-IFN-α or -β neutralizing antibodies, a polyclonal rabbit IgG antibody served as the control. For single i.p. injection, drugs were injected on d11 after tumor inoculation. For twice i.p. injection, drugs were delivered on d3 and d7 or d10 after LLC implantation, as detailed in the results and/or figure legends.

### Behavior tests

All the behavioral tests were conducted in a blinded manner and performed during between the hours of 9:00-16:00. Animals were habituated in a light and humidity controlled testing environment for at least 2 days prior to baseline testing. For Von Frey testing, mice were confined to individual 5×5 cm boxes placed on an elevated wire grid. A blinded experimenter stimulated their hindpaws using a series of von Frey filaments with logarithmically increasing stiffness (0.02– 2.56g, Stoelting). Each filament was applied perpendicularly to the central plantar surface. The 50% paw withdrawal threshold was determined using Dixon’s up-down method ^48^. In addition, we also measured paw withdrawal frequency to repeated stimulation (10 times, with ∼1-2 minutes between each stimulation) using a subthreshold 0.16g von Frey filament, which is a more sensitive method to detect mechanical allodynia. To measure cold allodynia, mice were similarly isolated to individual boxes on an elevated mesh floor, and a drop (∼20-30 µl) of acetone was applied to the plantar hindpaw. The duration of time that animal displayed a nociceptive response (lifting or licking the paw) over a 90s period immediately after acetone application was recorded. To measure locomotor function, we performed open field testing in which mice were placed in the center of a 45 x 45 cm chamber and locomotor activity was recorded by an overhead webcam connected to a laptop computer, and animals’ movements were automatically tracked for 30 minutes using ANY-Maze. The total distance traveled and mean speed during the 30-minute period were analyzed.

### *In vivo* X-ray radiography

Osteolytic bone destruction was continuously evaluated by radiography using the MultiFocus by Faxitron system (Faxitron Bioptics LLC, Tucson, Arizona). Radiographs of tumor-bearing femora were rated for bone destruction on a 0-5 score scale based on previous study ^27^: 0 for normal bone at baseline without tumor inoculation; 1 for one to three radiolucent lesions indicative of bone destruction compared to baseline; 2 for increased number of lesions (three to six lesions) and loss of medullary bone; 3 for loss of medullary bone and erosion of cortical bone; 4 for full-thickness unicortical bone loss; 5 for full-thickness bicortical bone loss and displaced skeletal fracture. All radiographic image quantifications were completed by an experimenter who was blinded to the experimental conditions.

### Microcomputed tomography

Microcomputed tomography (MicroCT) analyses were performed on femurs from tumor inoculated mice or naïve mice using a VivaCT 80 scanner with the 55-kVp source (Scanco, Southeastern, PA) as previously described ^27, 49, 50^. Quantification of microCT data was calculated for distal femurs of mice treated with vehicle or DMXAA. Parameters quantified included bone volume/total volume (BV/TV) and connectivity density (Conn.D) within a region of 100 slides and 200 slides proximal to the distal growth plate.

### Bone histology, TRAP, and ALP staining on mouse femurs

Mice were deeply anesthetized and perfused intracardially with 4% paraformaldehyde (PFA) in 0.1M phosphate buffered saline. The femora were removed and then post-fixed for 48h in the same fixative at 4 °C. After demineralization in EDTA (10%) for 10 days, femur samples were dehydrated in an ascending gradient of ethanol (30-100%) followed by paraffin embedding. Serial sections for trabecular bone were obtained from the distal femur at a thickness of 5 µm followed by tartrate-resistant acid phosphatase (TRAP) staining or alkaline phosphatase (ALP) staining using TRAP kits (Fast Red TR/Naphthol AS-MX, Sigma, St. Louis, MO) and NBT/BCIP (Thermo Scientific), respectively^27^. Bone static histomorphometric analyses for osteoclast number (osteoclast number per trabecular bone surface covered by osteoclasts, Oc.S/BS) and osteoblast number (osteoblast number per trabecular bone surface, Ob.N/BS) were conducted using Image J (NIH) based on images taken by a Leica Q500MC microscope. Osteoclasts, osteoblasts and trabecular bone at the metaphysis of the femur (1500 µm proximal to the distal growth plate) were quantified, since bone destruction in this model mainly occurs in this area ^33, 51^. Three sections per animal were randomly chosen and used for quantification.

### Visual and immunohistochemical analysis of lung metastases

Mice were deeply anesthetized with isoflurane and perfused intracardially with PBS, followed by 4% PFA. After the perfusion, lungs were removed from mice and post-fixed in the same fixative overnight. The samples were then dehydrated with a 30% sucrose solution, embedded in O.C.T. (Tissue Tek), and cryosectioned to produce 8 μm thick sections. For Hematoxylin and Eosin (H & E) staining, lung sections were rehydrated and stained with 0.1% Hematoxylin and 0.5% Eosin in sequence. After dehydration and clearance with HistoClear (Electron Microscopy Sciences, Hatfield, PA), slides were mounted with a resinous mounting medium (Mercedes Medical, Sarasota, FL) and subsequently imaged using a Leica Q500MC microscope with a digital camera at different magnifications.

### Bone marrow collection

Mice were humanely euthanized and femora were carefully removed and placed on ice. The muscles surrounding the femur were gently removed as much as possible, followed by separation of the distal epiphysis from femoral shaft. The proximal end of the femur removed, followed by insertion of a 25-gauge needle into the distal end of the femur and injection of 1 ml cold PBS or α-MEM (Gibco, Thermo Fisher Scientific) by lightly pressing down on the plunger, allowing the bone marrow to be evacuated into a 1.5 ml centrifuge tube. Bone marrow was harvested through centrifugation at 800g for 5 minutes at 4 °C for subsequent detections or cell cultures ^27^.

### ELISA

Mouse high-sensitivity IFN-α ELISA kit (42115-1) and IFN-β ELISA kit (42410-1) were purchased from PBL Assay Science. Mouse CTX-I ELISA kit (AC-06F1) and mouse PINP ELISA kit (AC-33F1) were purchased from Immunodiagnostic Systems. ELISA was performed using culture medium, serum, and bone marrow lysates. Serum was obtained from whole blood that was collected from a submandibular vein via facial vein puncture, coagulated for 30 minutes at room temperature, followed by centrifugation (2,000 × g for 10 min, 4°C) and collection of the supernatant (serum). Bone marrow was homogenized in a lysis buffer containing protease inhibitors at 4°C for 30 minutes. ELISA was conducted in accordance with the manufacturer’s instructions. A standard curve was performed for each experiment.

### *In vitro* induction of osteoclastogenesis and TRAP staining

For *in vitro* drug treatment, the RAW 264.7 cells were incubated with 35 ng/ml RANKL for six days. Isolated bone marrow cells were cultured overnight in α-MEM media containing 10% fetal bovine serum and 1% antibiotic-antimycotic solution. The suspended cells were collected and incubated with 20 ng/ml MCSF for 3 days to obtain bone marrow-derived macrophage (BMDM). The attached cells were further activated by 35 ng/ml RANKL and 20 ng/ml MCSF. Drug treatment was performed in tandem with RANKL. Culture medium and co-cultured reagents were changed every 3 days. After 6 or 7 days of incubation, the cells were fixed by 4% PFA and stained with warm TRAP staining solution (TRAP kit, Sigma-Aldrich, SLBW4002) for 10-30 min at 37°C^27^. TRAP-positive multinucleated cells that displayed three or more nuclei under a light microscope were considered osteoclasts, and the numbers of positive cells were counted in a blinded fashion with images of randomly selected visual fields (4-5 regions per well) using Image J software.

### Flow cytometry

For analysis of immune subsets present within the bone marrow tumor microenvironment, bone marrow was collected as previously described followed by removal of RBC cells using RBC lysis buffer (Sigma, R7757). Cells were subsequently washed with PBS and resuspended in 2% paraformaldehyde in PBS for 10 minutes. Fixed cells were washed several times with PBS, followed by incubation in blocking buffer (1% anti-mouse-CD16/CD32, 2.4 G2, 2% FBS, 5% NRS,1% triton x100 and 2% NMS in HBSS; BD Bioscience) for 1h at room temperature. Cells were subsequently stained with IL-17-FITC (1:20, rat, Miltenyi Biotec, 130-102-262), CD-3 APC/cy7 (1:200, rat, Biolegend, 100221), CD-4 APC (1:200, rat, Biolegend, 100411), FoxP3-PE (1:20, human, Miltenyl Biotec, 130-111-678), CD8a-FITC (1:200, rat, Biolegend, 100705), and CD11b-PE(1:200, rat, Biolegend, 101207) in blocking buffer for 1h at room temperature. After staining, cells were washed in PBS with EDTA. Flow cytometry events were acquired by a BD FACS Canto II flow cytometer using the BD FACS Diva 8 software (BD Bioscience). Data were analyzed using Cytobank software (https://www.cytobank.org/cytobank).

### *In vivo* bioluminescence imaging

RediJect D-Luciferin Ultra was purchased from PerkinElmer (770505). Prior to *in vivo* imaging, mice were shaved in the region of interest depicted in the figure. Bioluminescence images of LL/2- Luc2 bearing mice were captured with IVIS Lumina III system 15 min after intraperitoneal injection of D-Luciferin (30 mg/kg)^27^. The IVIS acquisition control panel was set to following conditions for imaging: Exposure time = auto, Binning = medium, F/Stop = 1, Emission Filter = open. The bioluminescence images were analyzed using Living Image software from PerkinElmer.

### Whole-cell patch clamp recordings in whole-mount DRGs ex vivo

Four-week-old male C57BL/6 mice were used to establish the bone cancer pain model by intrafemoral inoculation of LLC, leading to nociceptor hyperexcitability in the ipsilateral L3-L5 DRGs which extend afferent nerve fibers to the tumor-bearing femur. Notably, young mice were used for these experiments due to technical limitations in performing electrophysiological recordings on older mice. 11d after tumor implantation, mice were euthanized followed by careful isolation of L3-L5 DRGs, which were placed in oxygenated artificial cerebrospinal fluid. DRGs were lightly digested for 20 minutes using an enzymatic mixture consisting of 0.32 ml collagenase A (1 mg/mL) and Trypsin (0.25%). Intact DRGs were then incubated in ACSF oxygenated with 95% O2 and 5% CO2, supplemented with vehicle (PBS) or 30 µM DMXAA in PBS for 2 hours at 37°C. Following incubation, DRGs were transferred to a recording chamber, where neurons could be visualized using a 40x water-immersion objective on an Olympus BX51WI microscope. Patch pipettes were pulled from borosilicate capillaries (Chase Scientific Glass Inc.) and filled with a pipette solution containing (in mM): 126 potassium gluconate, 10 NaCl, 1 MgCl2, 10 EGTA, 2 Na-ATP, and 0.1 Mg-GTP, adjusted to pH 7.3 with KOH. The external solution was composed of (in mM): 140 NaCl, 5 KCl, 2 CaCl2, 1 MgCl2, 10 HEPES, 10 glucose, adjusted to pH 7.4 with NaOH. The resistance of pipettes was 4-5 MΩ. Series resistance was compensated for (>80%) and leak subtraction was performed. Data were low-pass filtered at 2 KHz and sampled at 10 KHz. Data were recorded and analyzed using the pClamp10 (Axon Instruments) software.

### Data analysis and statistics

The sample sizes for each experiment were based on our previous studies on such experiments^27, 45^. Statistical analysis was performed with GraphPad Prism 6.0 (GraphPad Software). All the data in the figures are expressed as mean ± standard error (SEM). Biochemical and behavioral data were analyzed using a two-tailed t-test (two groups), One-Way ANOVA, or Two-Way ANOVA, followed by post-hoc Bonferroni test. Fisher’s exact test was utilized for the comparison of the bone fracture ratio. The criterion of *P* < 0.05 was defined as the threshold for statistical significance. In each figure, significance denotes *P* values as follows: * *P* < 0.05; ** *P* < 0.01; *** *P* < 0.001, **** *P* < 0.0001.

### Study approval

The present studies in animals were reviewed and approved by the IACUCs of Duke University. All animal procedures were conducted in accordance with the NIH’s Guide for the Care and Use of Laboratory Animals (National Academies Press, 2011)

## Acknowledgements

We thank Dr. Duncan Lascelles for helpful discussions. This study was supported in part by Duke University Anesthesiology Research funds and Duke Microbiome Center Grant awarded to R-R.J.

## Author contributions

K.W., C.R.D. and R-R.J. designed experiments. K.W., C.R.D., C.J., X.T., S.B., and M.L. conducted behavioral, radiograph, ELISA, tissue culture, flow cytometry, histochemical, and electrophysiology experiments and analyzed the data. Y.L. conducted micro-CT experiments under supervision of M.J.H. K.W., C.R.D. and R-R.J. wrote the paper. M.J.H. edited the manuscript.

## Ethics declarations

The authors declare no competing interests. Dr. Ji is a consultant of Boston Scientific and received research support from Boston Scientific. These activities are not related to this study.

---

**Extended Data Fig. 1.**
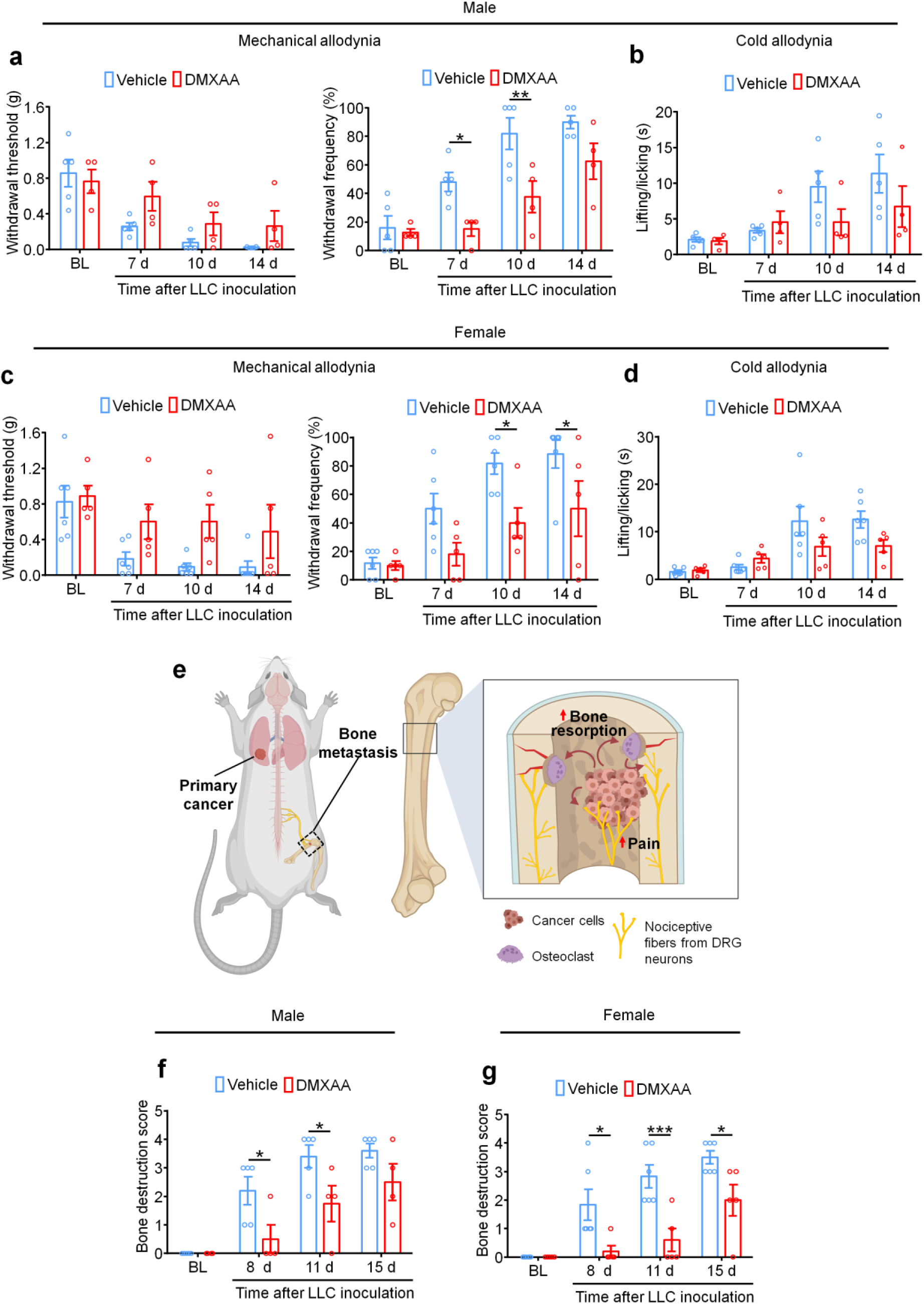
STING agonists reduce bone cancer pain and bone destruction in both male and female mice. **a-d.** Von Frey testing to determine mechanical allodynia (**a, c**) or acetone testing to determine cold allodynia (**b, d**) in male mice (**a, b**) and female mice (**c, d**), displayed separately. **e**. Schematic showing the mechanisms of bone cancer-induced pain, which originates from direct activation of local nociceptive afferents by mediators produced from cancer cells and osteoclasts as well as indirect mechanisms owed to osteoclast-induced bone resorption leading to subsequent bony fractures and breaks. **f-g**. Radiographic measurement of bone destruction following vehicle or DMXAA (2 x 20 mg/kg, i.p.) treatment at the indicated timepoints in male (**f**) or female mice (**g**). Sample sizes are indicated by individual data points (n = 4-5 mice/group). Data displayed are the mean ± SEM. \**P* < 0.05, \*\**P* < 0.01 and \*\*\**P* < 0.001, repeated-measures two-way ANOVA with Bonferroni’s *post-hoc* test.

**Extended Data Fig. 2.**
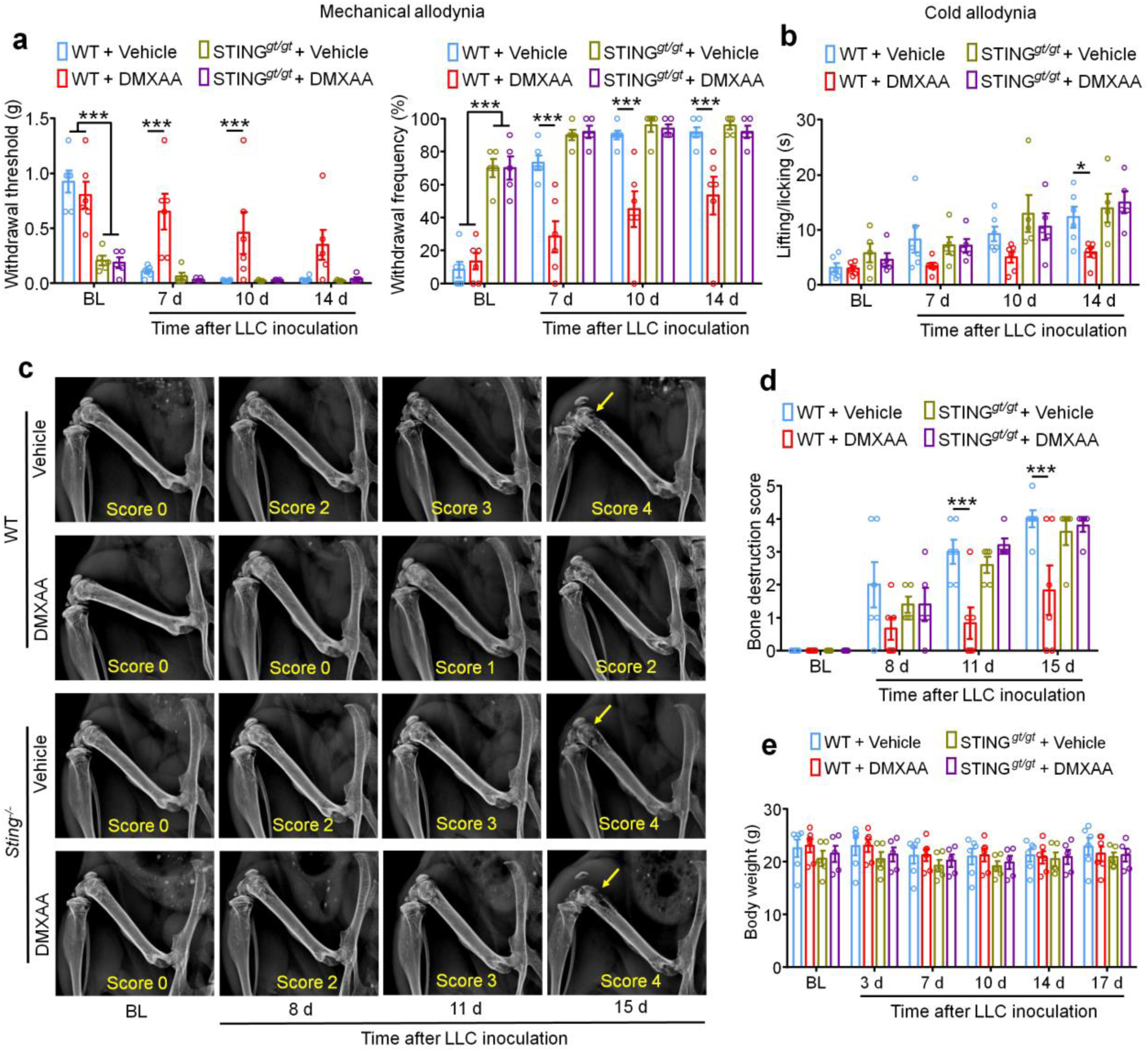
The protective effects of DMXAA treatment are STING dependent. **a.** Mechanical allodynia from von Frey tests in STING*^+/+^* (WT) and STING*^gt/gt^* mice treated with vehicle or DMXAA (2×20 mg/kg, i.p.). Left, withdraw threshold. Right, withdrawal frequency (n = 5-6 mice/group). **b.** Cold allodynia measured by acetone testing in STING*^+/+^* (WT) mice and STING*^gt/gt^* mice with the indicated treatments (n = 5-6). **c-d.** Radiography to measure bone destruction score at the indicated timepoints after tumor inoculation in STING*^+/+^* (WT) and STING*^gt/gt^* mice with vehicle or DMXAA treatment. (**c**) Representative radiographs of tumor-bearing femora. Bone destruction score is indicated in each image and arrows show bone lesions with scores over 3. (**d**) Quantification for (**c**) (n = 5-6 mice/group). **e.** Measurement of body weight after LLC inoculation in mice at the indicated timepoints and treatment groups (n = 5-6 mice/group). Data represent the mean ± SEM. \**P* < 0.05 and \*\*\**P* < 0.001, repeated-measures two-way ANOVA with Bonferroni’s *post-hoc* test.

**Extended Data Fig. 3.**
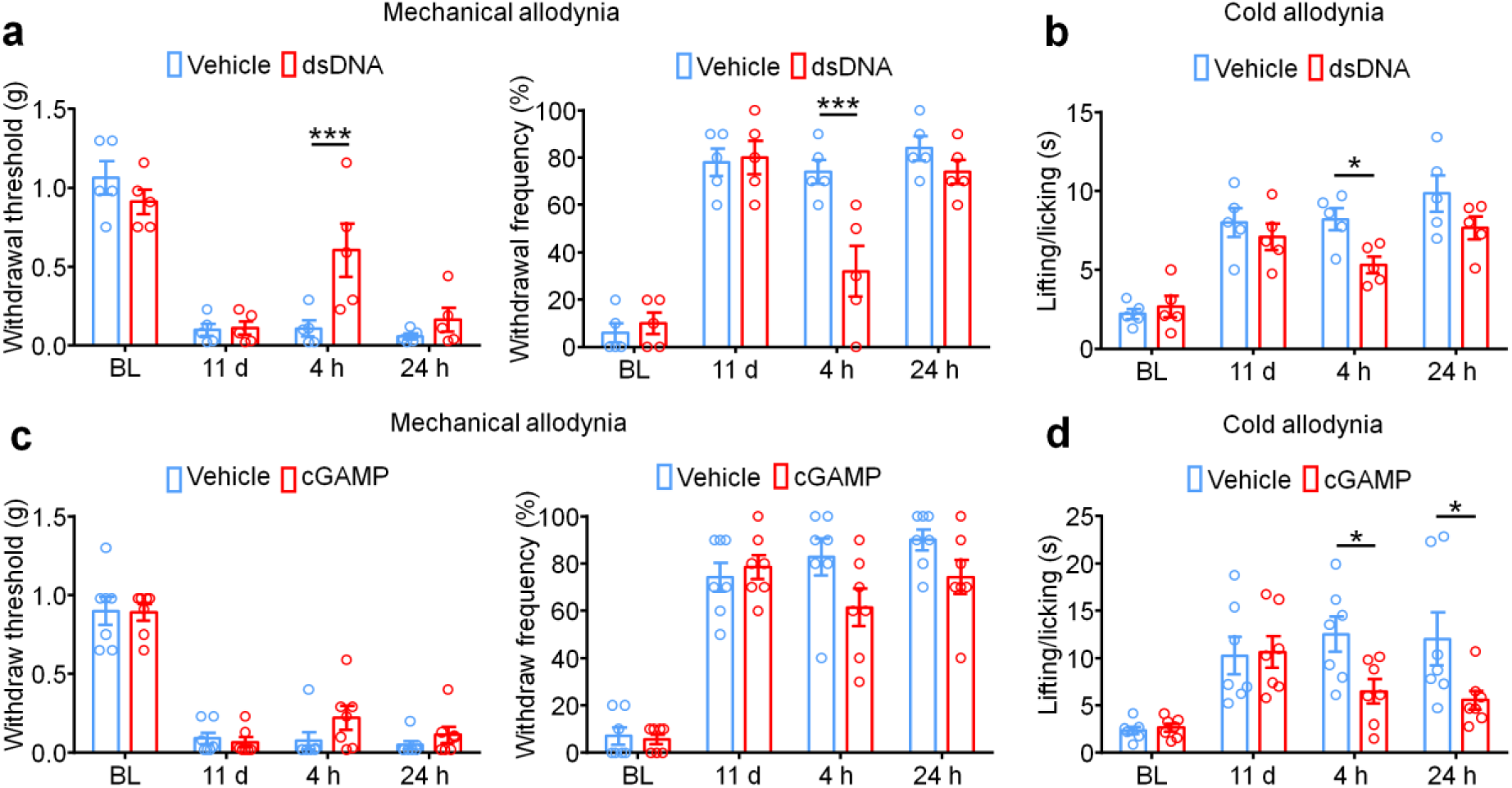
The natural STING activators dsDNA and cGAMP also suppress bone cancer pain. **a-b.** Measurement of cancer-induced mechanical allodynia (**a**) at d11 post LLC inoculation, as determined by withdrawal threshold (**a**, left) or withdrawal frequency (**a**, right) or cold allodynia as determined by the acetone test before or 4h or 24h after administration of the natural STING agonist dsDNA (30 µg, i.p., n = 5 mice/group). **c-d.** Mechanical allodynia measured by von Frey testing (**c**) or cold allodynia measured by acetone testing (**d**) before and 4h or 24h after treatment with vehicle or the natural STING ligand 3’3’-cGAMP (20 mg/kg, i.p.) on d11 after LLC inoculation (n = 7 mice/group). Data indicate the mean ± the SEM. \**P* < 0.05, repeated-measures two-way ANOVA with Bonferroni’s *post hoc* test. Data are Mean ± SEM. \**P* < 0.05 and \*\*\**P* < 0.001, repeated-measures two-way ANOVA with Bonferroni’s *post-hoc* test.

**Extended Data Fig. 4.**
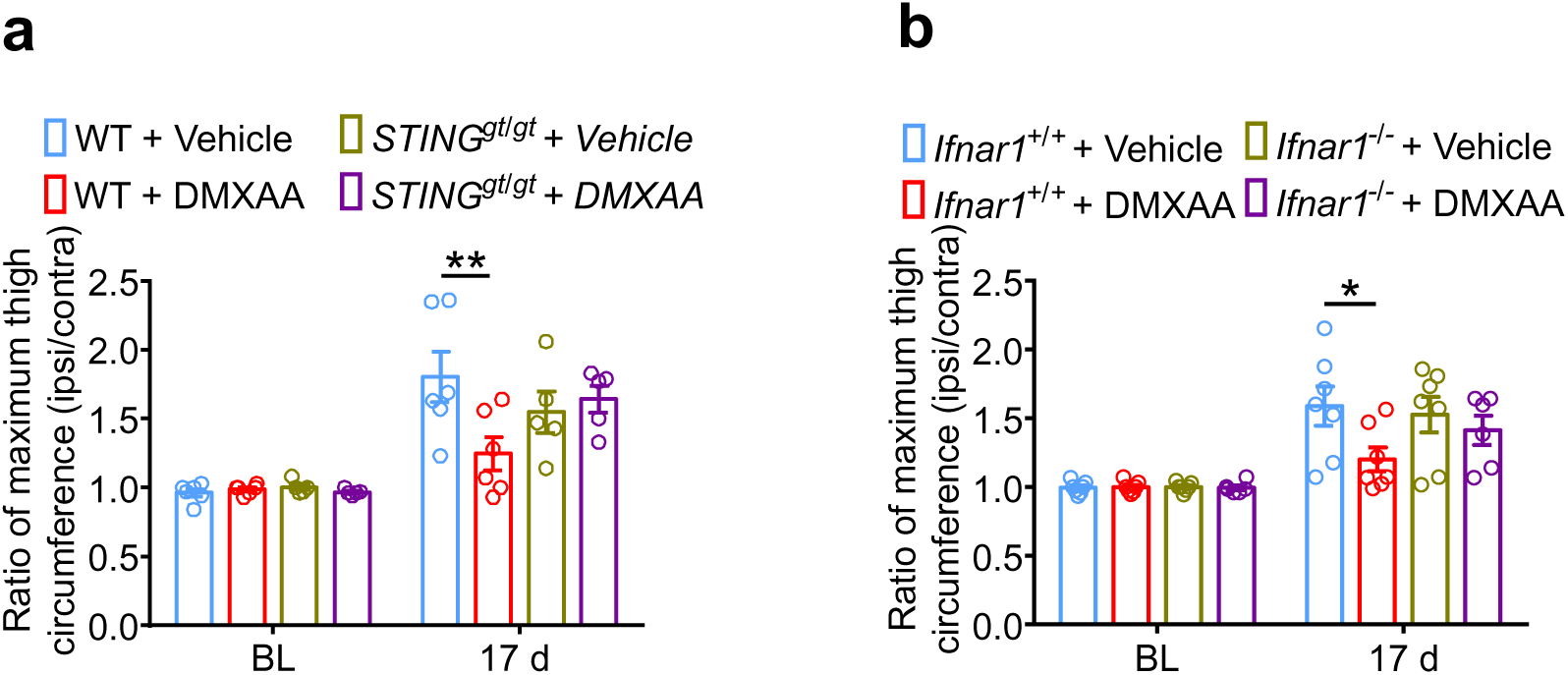
Reduction in local tumor burden by DMXAA requires host-intrinsic STING and IFN-I signaling. **a.** Ratio of maximum thigh circumference in WT mice or STING*^gt/gt^* mice treated with vehicle or DMXAA (2 x 20 mg/kg, i.p.) on day 17 after LLC implantation, n = 5-6. **b.** Ratio of maximum thigh circumference in WT mice or STING*^gt/gt^* mice treated with vehicle or DMXAA (2 x 20 mg/kg, i.p.) on day 17 after LLC implantation, n = 6-7. Data are Mean ± SEM. \**P* < 0.05, repeated-measures two-way ANOVA with Bonferroni’s *post hoc* test.

**Extended Data Fig. 5.**
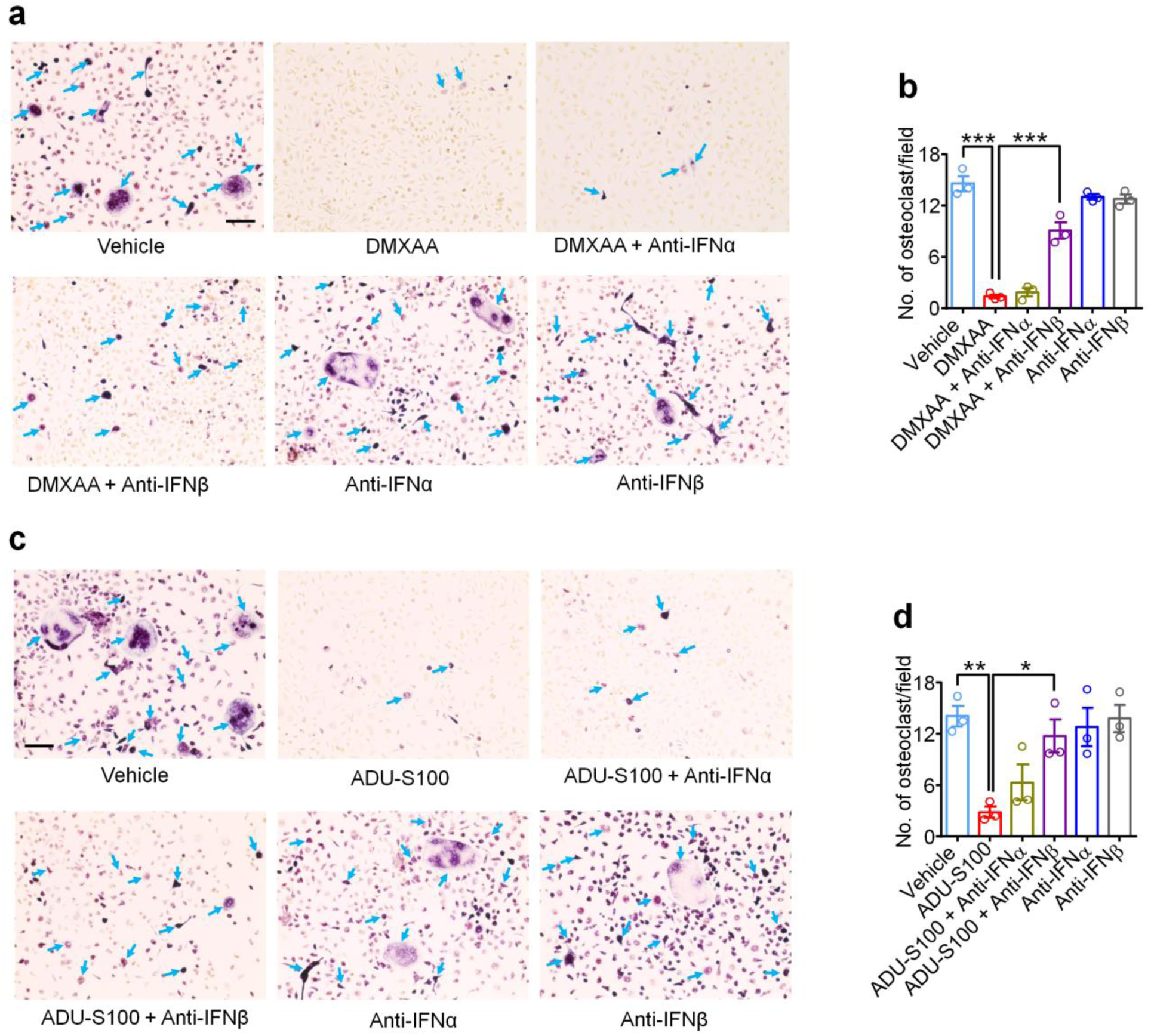
STING agonists suppress osteoclastogenesis via IFN-α and IFN-β. **a-d.** TRAP staining after treatment of DMXAA **(a, b)** and ADU-S100 **(c, d).** (**a, c**) Representative images of TRAP staining to identify *in vitro* BMDM-derived osteoclasts following differentiation with RANKL (35 ng/ml) and M-CSF (20 ng/ml), together with treatment of DMXAA (30 µM) and/or anti-IFN-α antibody (600 ng/ml) or anti-IFN-β antibody (600 ng/ml) (**a**) or treatment of ADU-S100 (30 µM) and/or anti-IFN-α antibody (600 ng/ml) or anti-IFN-β antibody (600 ng/ml) (**c**), Scale bar, 100 µm. (**b**, **d**) Quantification for left (n = 3). Data displayed represent the mean ± SEM. \*\*\**P* < 0.001, one-way ANOVA with Bonferroni’s *post-hoc* test.

**Extended Data Fig. 6.**
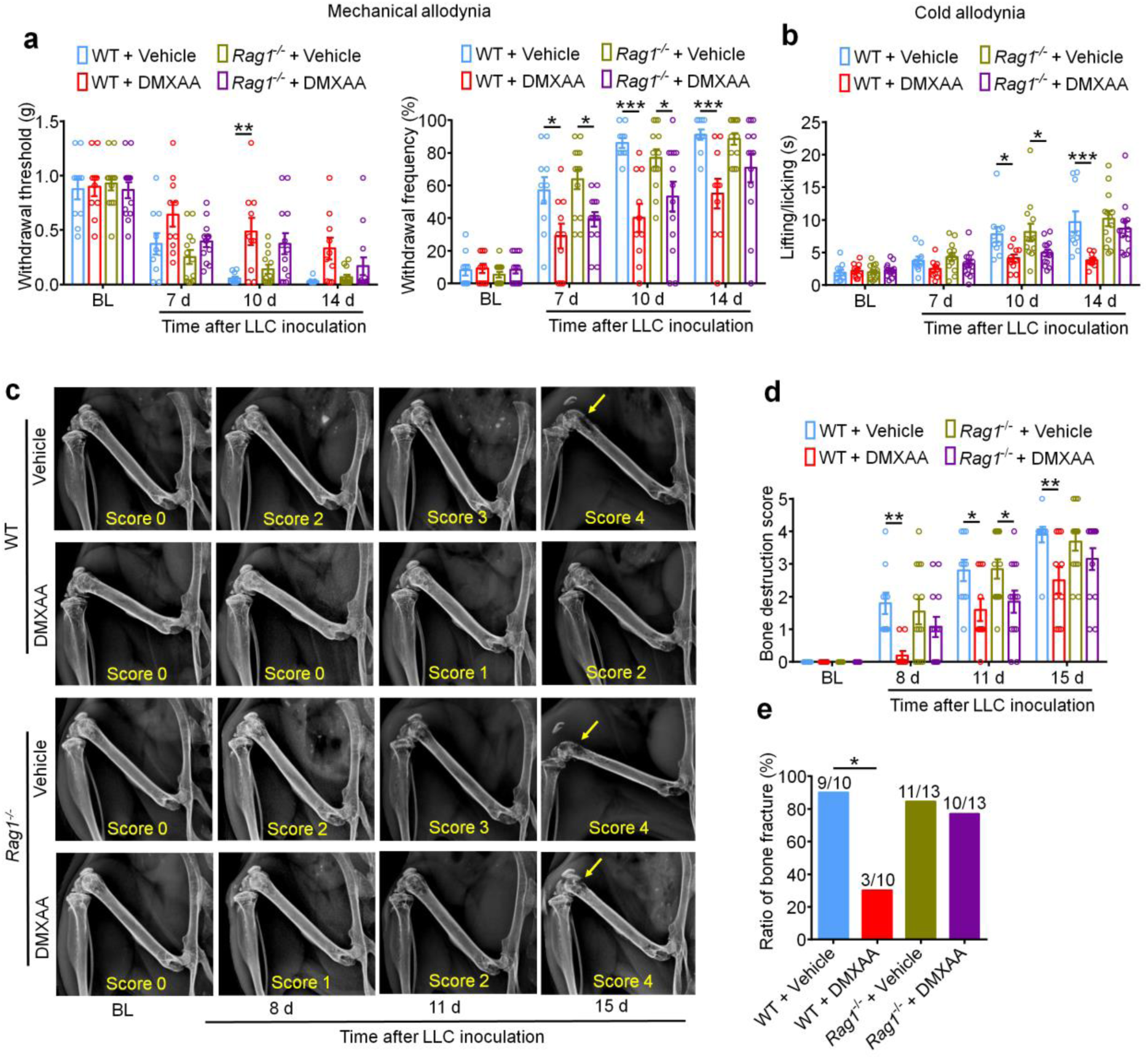
The analgesic and bone-protective effects of STING agonists remain intact until late stages in T cell-deficient *Rag1^−/−^* mice. **a.** Mechanical allodynia from von Frey test in WT or *Rag1^−/−^* mice treated with vehicle or DMXAA (2 x 20 mg/kg, i.p.) on baseline (BL), day 7, 10 and 14 after tumor inoculation (n = 10-13). Left, withdrawal threshold. Right, withdrawal frequency. **b.** Cold allodynia from acetone test in WT or *Rag1^−/−^* mice with indicated therapy (n = 10-13). **c-d.** Radiographical analysis of bone destruction in WT or *Rag1^-/-^* mice administered vehicle or DMXAA, measured at BL, d8, d11 and d15 post LLC inoculation. (**c**) Representative X-ray images. Bone destruction score is labeled on the bottom of each photo and arrows indicate bone destruction scores of more than 3. (**d**) Quantification for (b) (n = 10-13). **e.** Quantification of the proportion of mice with distal bone fractures, harvested and analyzed at d17 post-inoculation and with the indicated genotypes and treatment groups. Data are Mean ± SEM. \**P* < 0.05, \*\**P* < 0.01 and \*\*\**P* < 0.001, repeated-measures two-way ANOVA with Bonferroni’s *post hoc* test.

**Extended Data Fig. 7.**
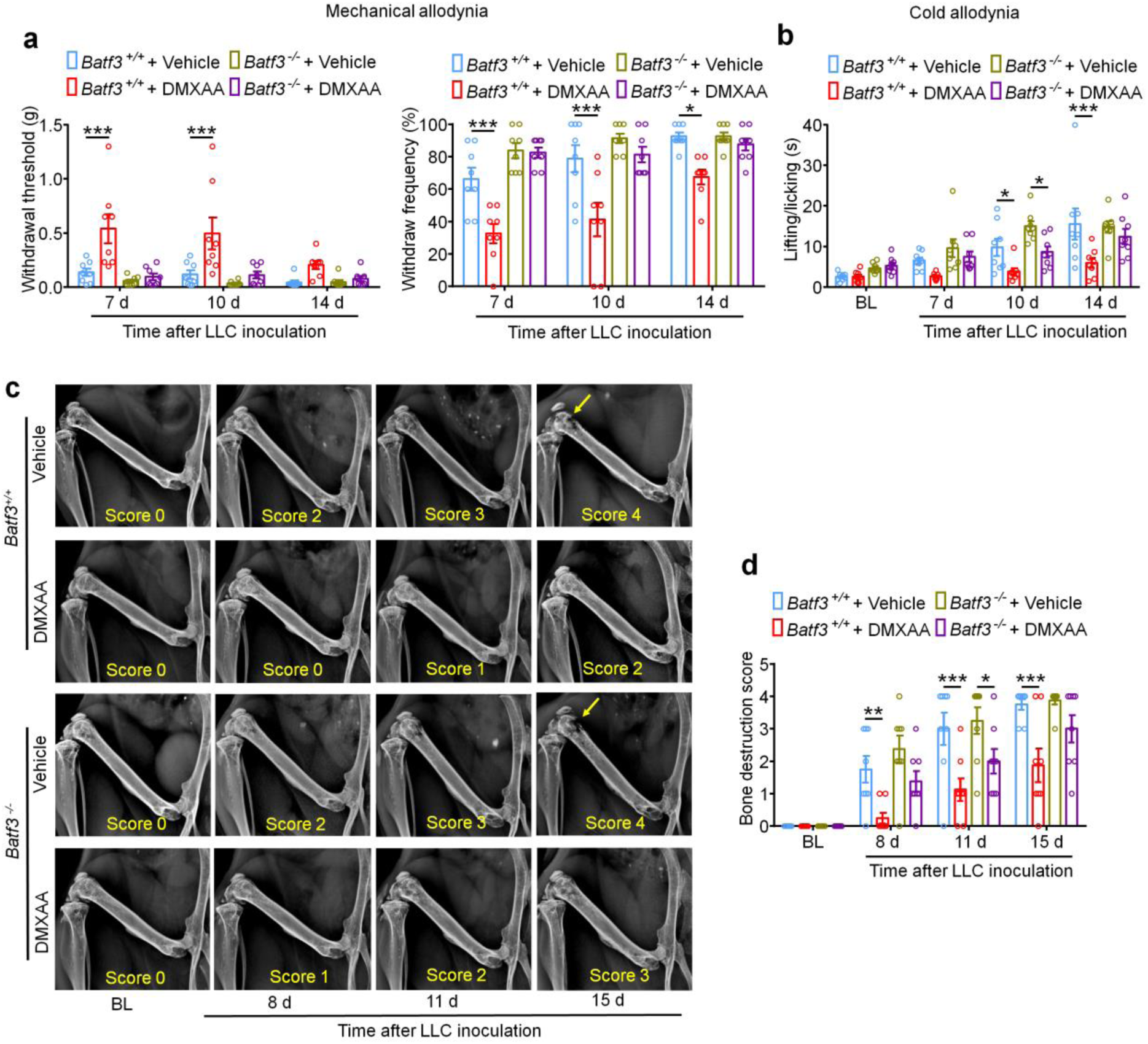
The protective effects of STING agonists remain largely intact until late stages in *Batf3^−/−^* mice. **a.** Mechanical allodynia from von Frey test in *Batf3^+/+^* or *Batf3^−/−^* mice treated with vehicle or DMXAA (2 x 20 mg/kg, i.p.) on baseline (BL), day 7, 10 and 14 after tumor inoculation (n = 8). Left, withdrawal threshold. Right, withdrawal frequency. **b.** Cold allodynia from acetone test in *Batf3^+/+^* or *Batf3^−/−^* mice with indicated therapy (n = 8). **c-d.** Radiographical analysis of bone destruction in *Batf3^+/+^* or *Batf3^−/−^* mice administered vehicle or DMXAA, measured at BL, d8, d11 and d15 post LLC inoculation. (**c**) Representative X-ray images. Bone destruction score is labeled on the bottom of each photo and arrow indicates bone destruction score more than 3. (**d**) Quantification for (**c**) (n = 8). Data are Mean ± SEM. \**P* < 0.05, \*\**P* < 0.01 and \*\*\**P* < 0.001, repeated-measures two-way ANOVA with Bonferroni’s *post hoc* test.

**Extended Data Table 1.**
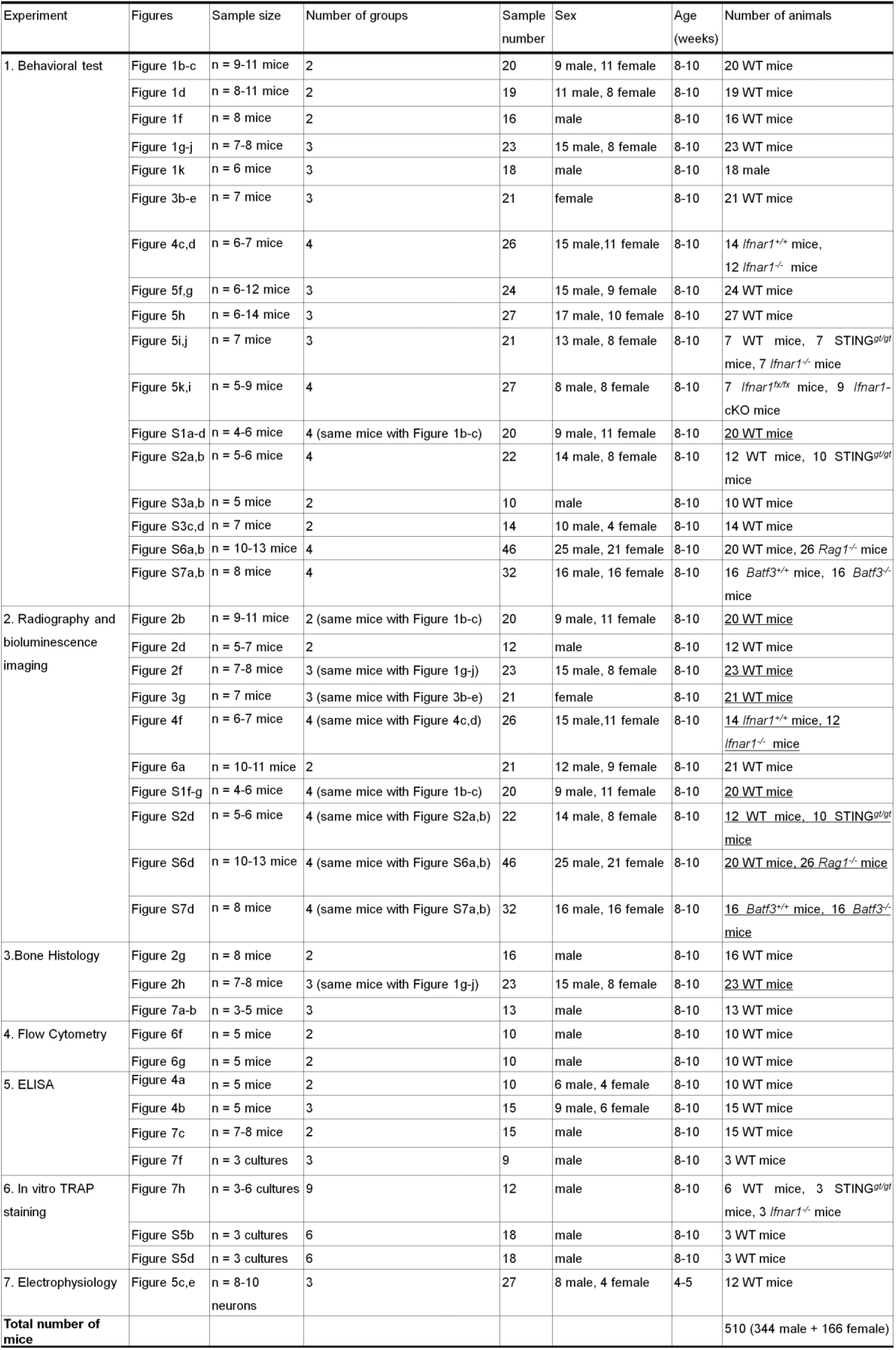
Number of animals used in different experiments. Both sexes of mice were used in the present study. Mice shared in different experiments are underlined. A total of 510 mice were used, including 344 males and 166 females.

